# Engineering a pacemaker-driven human mini-heart guided by spatial and single cell multi-omics of sinoatrial node development

**DOI:** 10.64898/2026.05.07.723626

**Authors:** Jiajun Zhu, Zhaowei Han, Roberto De Gregorio, Kevin Chang, Zheyu Zhang, Xue Dong, Kalpita Banerjee, Kai Liu, Martha Rea-Moreno, Mikal Kizilbash, Alicia Alonso, Jie Liu, Su-Yi Tsai, Ya-Wen Chen, Todd Evans, Shuibing Chen

**Affiliations:** Department of Surgery, Weill Cornell Medicine, 1300 York Ave, New York, NY, 10065, USA; Center for Genomic Health, Weill Cornell Medicine, 1300 York Ave, New York, NY, 10065, USA; Department of Computational Medicine & Bioinformatics, University of Michigan, Ann Arbor, MI, 48109, USA; Caryl and Israel Englander Institute for Precision Medicine, Weill Medical College of Cornell University, New York, NY 10065, USA; Epigenomics Core Facility, Weill Cornell Medicine, New York, NY 10065, USA; Department of Otolaryngology, Department of Cell, Developmental and Regenerative Biology, Institute for Airway Sciences, Institute for Regenerative Medicine, Icahn School of Medicine, New York, NY 10029, USA; Department of Life Science, National Taiwan University, Taipei 10617, Taiwan

## Abstract

The human sinoatrial node (SAN) functions as the primary pacemaker of the heart and coordinates the hierarchical electrical activity that drives cardiac contraction. However, experimental systems capable of reconstructing pacemaker-driven cardiac organization in human tissues remain limited. Here we integrate spatial multi-omics of the human fetal SAN with stem-cell engineering to generate pacemaker organoids (“Sinoids”) and assemble them into a pacemaker-driven human mini-heart composed of sinoatrial, atrial and ventricular cardiac modules. High-resolution spatial transcriptomics and single-nucleus multi-omic analyses of human fetal SAN tissues identify regulatory pathways guiding pacemaker lineage specification, which we leverage to engineer human pluripotent stem cell–derived SAN organoids with robust pacemaker identity and electrophysiological activity. When integrated with atrial and ventricular cardioids, Sinoids initiate and coordinate electrical activation across assembled cardiac tissues, establishing directional propagation of electrophysiological signals within structured mini-heart organoids. Combining AI-guided perturbation modeling with functional validation further identifies conserved regulatory pathways controlling pacemaker specification and regionalization, including YAP–TEAD and NRG–ERBB signaling. Together, these results establish a multi-omic–guided strategy for engineering pacemaker tissues and reconstructing cardiac conduction hierarchy in vitro. The pacemaker-driven mini-heart platform provides a modular human cardiac system for studying pacemaker biology, modeling arrhythmia mechanisms and enabling electrophysiological drug discovery.

## Main

The human sinoatrial node (SAN) functions as the dominant pacemaker of the heart, autonomously generating rhythmic electrical impulses that initiate each cardiac cycle and coordinate electromechanical coupling across cardiac chambers^1,2^. Despite its essential physiological role, the mechanisms governing pacemaker lineage specification, regional heterogeneity within the SAN, and functional integration with working myocardium in humans remain incompletely understood. Single-cell and single-nucleus multi-omics (sn-multiomics) data have enabled high-resolution interrogation of human heart development. scRNA-seq studies have charted early cardiac progenitors^3–5^ and cardiac chambers^3,6,7^, revealing extensive cellular diversity across fetal and adult human hearts^8–20^ and complementary murine systems^21^. Application of microdissection-coupled scRNA-seq has further resolved the cardiac conduction system (CCS) at single-cell resolution^6^. Spatial transcriptomics has mapped transcriptional programs within intact tissue architecture, illuminating spatially organized regulatory domains that govern chamber specification and septation^8,22–24^. Together, these atlases provide an increasingly detailed descriptive framework of cardiac cell states.

However, descriptive atlases alone do not establish regulatory causality, nor do they readily translate into experimentally tractable systems for reconstructing human pacemaker tissues. A key challenge remains bridging atlas-level annotations with mechanistic reconstruction of lineage decisions and tissue organization *in vitro*. Integrating developmental multiomics with stem cell engineering offers a promising strategy to address this gap, enabling insights from human developmental datasets to guide the design of functional tissue models. Here we integrate multiomic profiling of the human fetal SAN with functional perturbation and stem cell–derived tissue engineering to define a stage-resolved regulatory architecture underlying human pacemaker specification. We further test whether the inferred regulatory logic can guide the generation of stable pacemaker tissues and support the assembly of engineered cardiac organoids with functional pacemaker activity, thereby linking developmental genomics with human tissue engineering.

### Single-nucleus multiomics (sn-multiomics) analysis of the human fetal SAN

To resolve the cellular composition and regulatory landscape of the human SAN at single-cell resolution, we first performed sn-multiomics analysis, using paired single-nucleus RNA sequencing (snRNA-seq) and single-nucleus assay for transposase-accessible chromatin sequencing (snATAC-seq), on four human fetal heart samples that included the SAN region collected between 14- and 17-weeks post conception (**Table S1**). Following stringent quality control (QC; Methods), we retained 28,892 high-quality nuclei with paired snRNA-seq and snATAC-seq data for downstream analyses.

Unsupervised clustering identified 13 major cell clusters with distinct transcriptional profiles and corresponding chromatin accessibility patterns (**Fig. 1a**). These identities were further confirmed using weighted nearest neighbor (WNN) analysis, which integrates multimodal data to achieve optimal clustering resolution (**Fig. 1b**). The identified clusters include *NPPA^+^* atrial cardiomyocytes (ACM), *CDH5*^+^ endothelial cells, *WT1*^+^ epicardial cells, *PAX1*^+^ epithelial cells, *PDGFR*A^+^ fibroblasts, *CD96*^+^ lymphoid cells, *CD163*^+^ myeloid cells, *SYN3*^+^ or *NRXN1*^+^ neuronal cells, *HBG1*^+^ red blood cells (RBC), *HCN1*^+^ SAN cells, *MYH11*^+^ vascular smooth muscle (SMC), and *MYL2*^+^ ventricular cardiomyocytes (VCM). RNA-level expression and chromatin accessibility of key marker genes validated the identity of these populations (**Fig. 1c**, **1d** and **Extended Data Fig. 1a-1c**). SAN cells were enriched for canonical pacemaker markers, including *SHOX2*, *HCN4*, *HCN1*, *CACNA1G*, *CACNA2D2* and *GJC1* (**Fig. 1e**).

**Figure 1.**
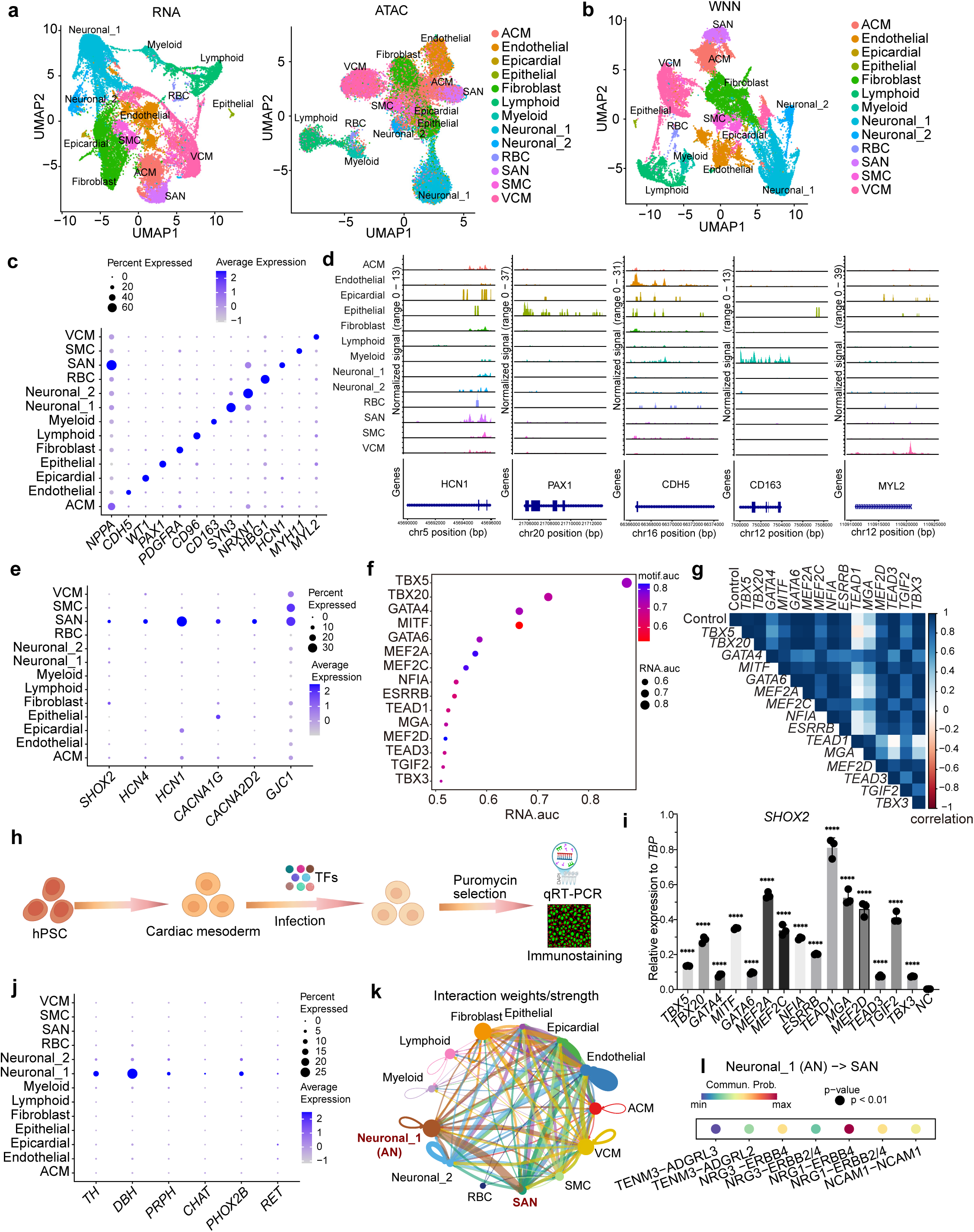
Integrated sn-multiomics profiling of human fetal SAN samples. **a,** UMAP plots of snRNA-seq (left) and snATAC-seq (right) showing clustering of 28,892 nuclei from human fetal SAN tissues. **b,** UMAP plot of weighted nearest neighbor (WNN) integration highlighting major cell types based on combined transcriptomic and chromatin accessibility profiles. **c,** Dot plot illustrating RNA-level expression of representative marker genes across identified cell populations. **d,** IGV tracks illustrating chromatin accessibility at selected cell type–specific loci. **e,** Dot plot illustrating expression of SAN-specific gene expression. **f,** Motif enrichment analysis ranked by average motif AUC and RNA AUC, identifying candidate TFs associated with SAN identity. **g,** Correlation analysis comparing control and individual perturbation targets based on predicted SAN marker gene expression following AI-facilitated virtual perturbation. **h,** Schematic overview of the experimental workflow used to evaluate regulatory TFs. **i,** qRT-PCR analysis of *SHOX2* expression in Day 20 cells following TF overexpression. Data are shown as mean ± SEM; p**** < 0.0001; N = 3 biological replicates. **j,** Dot plot illustrating AN gene expression. **k,** CellChat network plot depicting inferred intercellular signaling interactions among all identified cell populations. **l,** CellChat analysis illustrating predicted Neuronal_1(AN)-to-SAN signaling interactions.

We next leveraged the paired transcriptomic and epigenomic data to identify key transcription factors (TFs) involved in SAN development. We prioritized TFs that were both enriched at the RNA level in SAN cells and exhibited increased chromatin accessibility at their corresponding binding motifs, as assessed using the *chromVAR* package^25^. This integrative approach identified 15 candidate TFs in SAN cells: *TBX5*^26^*, TBX20*^27^*, GATA4*^28,29^*, MITF, GATA6, MEF2A, MEF2C, NFIA, ESRRB, TEAD1, MGA, MEF2D, TEAD3, TGIF2,* and *TBX3* (**Fig. 1f**). To predict the effects of TF perturbations, we adopted the STATE architecture, a transformer-based model that represents perturbations using 5,120-dimensional ESM2 protein embeddings^30^ and predicts post-perturbation gene expression from control cell states. The model was trained on CRISPRi Perturb-seq datasets^30–32^. The snRNA-seq gene expression profiles from the sn-multiomics data were used as the control (unperturbed) cell population for the model to make predictions on. We subsampled 10,000 cells while preserving cell type proportions, and the model predicted how each cell transcriptome would change upon knockdown of each target TF. Using this framework, we found that, relative to control, perturbation of *TEAD1* and *MGA* produced the most pronounced effects on SAN transcriptional signatures (**Fig. 1g**). To experimentally validate these predictions, we overexpressed each candidate TF at the cardiac mesoderm stage, using lentiviral infection followed by puromycin selection and continued differentiation to day 20 (**Fig. 1h**). By day 20, spontaneous beating was observed exclusively in cultures overexpressing *TEAD1, MEF2A,* or *MGA*, but not under control conditions (**Extended Data Movies 1-4**). Consistent with the in silico predictions, qRT-PCR confirmed that all TFs, but most notably *TEAD1* significantly upregulated the pacemaker gene *SHOX2* (**Fig. 1i**).

Among the two neuronal populations identified, the Neuronal_1 cluster corresponded to autonomic neurons (ANs), expressing markers of both sympathetic neurons (*TH*, *DBH*, *PRPH*) and parasympathetic neurons (*PHOX2B*, *CHAT*, *RET*) (**Fig. 1j** and **Extended Data Fig. 2a**). Cell–cell communication analysis using CellChat revealed strong directional interactions from ANs to SAN cells (**Fig. 1k**, and **Extended Data Fig. 2b**), including signaling through the NRG1–ERBB4, NRG1–ERBB2/4, NRG3–ERBB4, NRG3–ERBB2/4, and NCAM1–NCAM1 ligand–receptor pairs (**Fig. 1l**). Among these pathways, neuregulin (NRG) signaling was the most prominent, with ANs exhibiting the strongest outgoing signaling strength and SAN cells showing the highest incoming signaling strength (**Extended Data Fig. 2b-2c**), highlighting a key role for AN-to-SAN NRG signaling during human SAN development.

**Figure 2.**
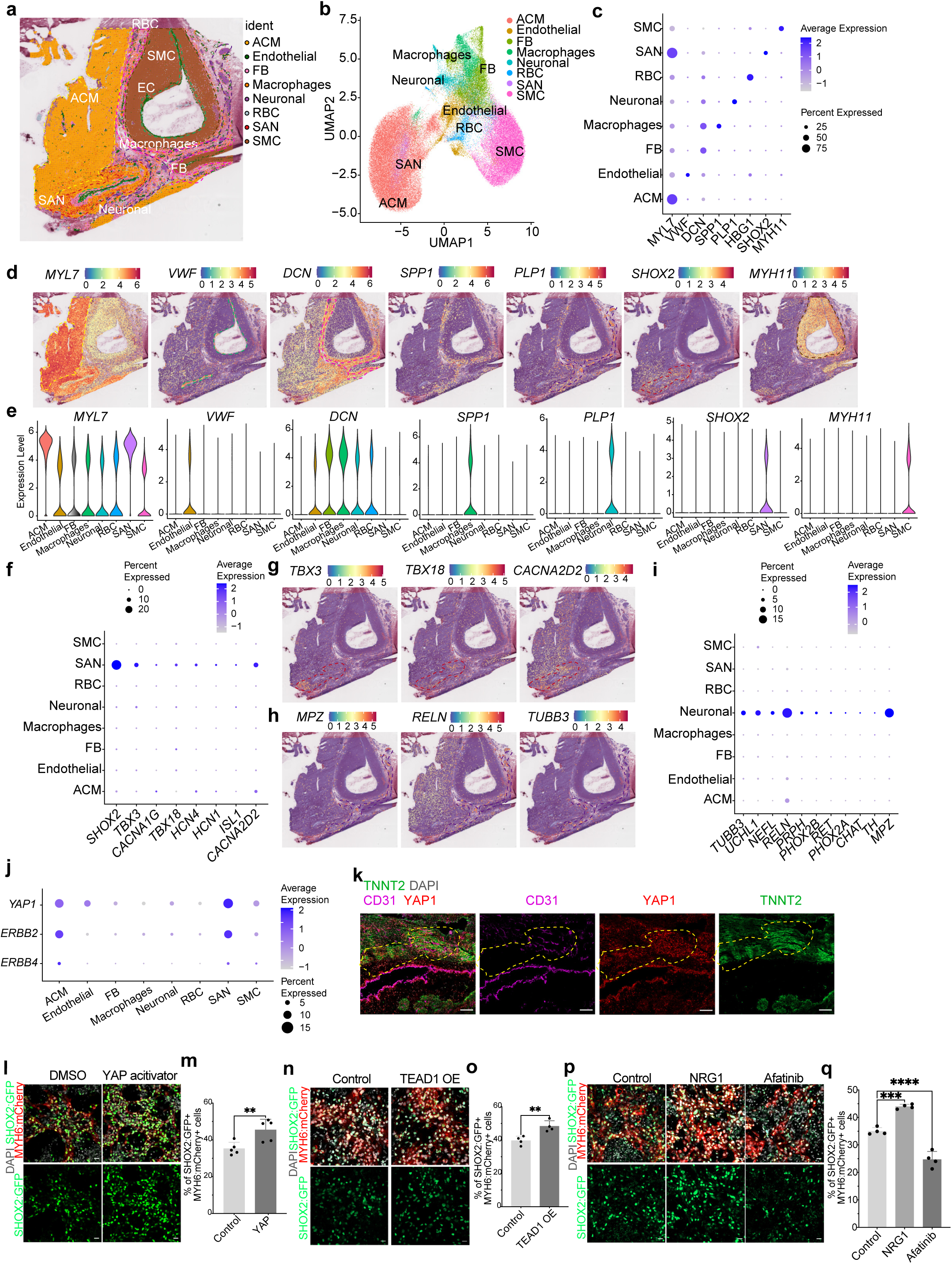
Spatial transcriptomic (Visium HD) analysis of human fetal SAN. **a**, Representative spatial image of human fetal SAN tissue section. **b**, UMAP plot illustrating the distribution of cell type specific markers. **c**, Dot plot illustrating the expression of representative marker genes. **d**, Spatial plots illustrating the localization of representative cell type specific markers. **e**, Violin plots illustrating the distribution of cell type–specific marker expression. **f**, Dot plot illustrating the expression of SAN-specific marker genes. **g**, Spatial localization of *TBX3*, *ISL1*, *TBX18* expression. **h**, Spatial localization of *MPZ*, *RELN*, *TUBB3* expression. **i**, Dot plot illustrating AN marker genes. **j**, Dot plot illustrating the expression of *ERBB2*, *ERBB4* and *YAP1*. **k**, Confocal images illustrating the expression of YAP1, CD31 and TNNT2 of SAN tissue slide. Scale bar=100 µm. **l**, **m,** Representative confocal images (I) and quantification of SHOX2:GFP+MYH6:mCherry+ cells (m) of day 20 hPSC-derived SAN after 5 uM kaempferol or DMSO treatment during differentiation. Scale bar=20 µm. Data are shown as mean ± SEM; **p < 0.01; N = 3 biological replicates. **n**, **o,** Representative confocal images (n) and quantification of SHOX2:GFP+MYH6:mCherry+ cells (o) of day 20 hPSC-derived SAN cells with control or *TEAD1* overexpression. Scale bar=20 µm. Data are shown as mean ± SEM; **p < 0.01; N = 3 biological replicates. **p**, **q,** Representative confocal images (p) and quantification of SHOX2:GFP^+^MYH6:mCherry^+^ cells (q) of day 20 hPSC-derived SAN cells following 50 ng/ml NRG1 or 5 µM afatinib treatment compared to control. Scale bar=20 µm. Data are shown as mean ± SEM; **p < 0.01; N = 3 biological replicates.

### Spatial transcriptional analysis of human fetal sinoatrial node (SAN)

We next performed High-Resolution (∼2 µm) spatial transcriptomic analysis of human fetal SAN tissue at a comparable developmental stage (17-weeks post conception). The tissue encompassing the superior region of the right atrium, where the SAN resides, was carefully microdissected, and the SAN location was confirmed by immunostaining for HCN4 (**Extended Data Fig. 3a**). Whole-transcriptome spatial profiling was performed using the 10x Genomics Visium HD 3’ platform. Unsupervised clustering identified eight major clusters, including *SHOX2^+^* SAN, *MYL7^+^* ACM^33^, *PLP1^+^* neuronal, *MYH11^+^*SMC, *SPP1^+^* macrophages^34^, *DCN^+^* fibroblasts (FB), *VWF^+^* endothelial cells and *HBG1^+^* RBCs (**Fig. 2a–2e**, **Extended Data Fig. 3b**). SAN also express other known markers of SAN, including *TBX3, CACNA1G*^35,36^*, TBX18, HCN4, HCN1, ISL1* and *CACNA2D2*^6,37^ (**Fig. 2f, 2g**).

Spatial mapping revealed that SAN cells are directly adjacent to both ACM and neurons, consistent with their biological role in regulating heart rhythm under the control of ANs^38^. The adjacent neuronal regions express general AN markers including *MPZ*, *RELN*, and *TUBB3* (**Fig. 2h**), as well as sympathetic neuron markers (*TH, DBH*) and parasympathetic neuron makers (*CHAT*, *PHOX2B*) (**Fig. 2i, Extended Data Fig. 3c**), suggesting they represent the Neuronal_1 subcluster identified by sn-RNA-seq (**Fig. 1a**). Consistent with the AI-facilitated TF analysis and Cell-Chat analysis in sn-multiomics data analysis, *YAP1,* the co-activator of *TEAD1*^39,40^, and *ERBB2, ERBB4*, the key receptors of NRG signaling, are highly expressed in the SAN region (**Fig. 2j**). Immunostaining further validates the expression of YAP in the SAN region of human fetal SAN tissue (**Fig. 2k**).

To functionally validate the roles of YAP/TEAD and NRG signaling in SAN specification, we used our previously established *SHOX2:EGFP; MYH6:mCherry* H9 hESC reporter line^41^ and differentiation protocol^42^ and modulated these signaling pathways. First, treatment with kaempferol, a YAP activator^43,44^, after mesoderm stage (day 3 to day 5) of differentiation (**Extended Data Fig. 4a**), significantly increased the proportion of SHOX2:EGFP⁺MYH6:mCherry⁺ SAN cells (**Fig. 2l, 2m**), accompanied by increased SHOX2:EGFP fluorescence intensity (**Extended Data Fig. 4b**). Consistent with this result, forced expression of *TEAD1* at the mesoderm stage (day 3) of differentiation (**Extended Data Fig. 4c**), similarly enhanced the percentage of SHOX2:EGFP⁺MYH6:mCherry⁺ SAN cells (**Fig. 2n, 2o**), as well as the intensity of SHOX2:EGFP expression (**Extended Data Fig. 4d**). Finally, treatment with NRG1 ligand at the mesoderm stage (day 3 to day 5, **Extended Data Fig. 4e**) markedly increased both the proportion of SHOX2:EGFP⁺MYH6:mCherry⁺ SAN cells and SHOX2:EGFP intensity, as assessed by immunostaining (**Fig. 2p**, **2q, Extended Data Fig. 4f**) and qRT-PCR (**Extended Data Fig. 4g**). In contrast, inhibition of ERBB2/4 receptor by afatinib produced the opposite effect (**Fig. 2q**).

To assess the *in vivo* relevance of YAP/TEAD and ERBB signaling in SAN development, we examined the effects of YAP and ERBB pathway inhibition in zebrafish embryos during specification of the pacemaker cells. For this purpose, we used the Tg(*nkx2.5*:*ZsYellow*) reporter strain^45^ and quantified pacemaker cell formation using immunofluorescence for Isl1. The zebrafish SAN-like structure is formed by Isl1^+^, Nkx2.5^-^ pacemaker cells that differentiate around the inflow tract at the base of the Nkx2.5^+^, Isl1^-^ atrial myocardium^46^. The primitive heart tube is formed by around 24 hours post fertilization (hpf) and the pacemaker cells can be imaged by around 48 hpf (**Extended Data Fig. 5a, 5b**). Compared with vehicle-treated controls, embryos treated starting at 30 hpf with afatinib (ERBB receptor antagonist) or BAY-593 (a YAP inhibitor)^47^ exhibited a marked reduction in the SAN cell population at 54 hpf (**Extended Data Fig. 5c**), as indicated by a diminished pool of Isl1^+^ cells (**Extended Data Fig. 5d**), as well as total Isl1^+^ area (**Extended Data Fig. 5e**). Drug treatments did not impact nkx2.5:ZsYellow expression from the heart tube or pharyngeal arches. Together, these results demonstrate the critical role of YAP/TEAD and NRG signaling in SAN lineage specification *in vivo*. Together, spatial, computational, *in vitro*, and *in vivo* analyses converge to demonstrate that YAP/TEAD and NRG–ERBB signaling act as conserved, functionally essential regulators of SAN lineage differentiation.

### hPSC-derived SAN organoid (Sinoid) as a model of SAN development

To more faithfully recapitulate the three-dimensional (3D) architecture and cellular complexity of the SAN, we developed an advanced differentiation strategy to generate 3D SAN organoids (Sinoids), incorporating stage-specific modulation of YAP and NRG1 signaling (**Extended Data Fig. 6**). As controls (**Extended Data Fig. 6**), we established parallel protocols to generate atrial cardioids (ACOs) and ventricular cardioids (VCOs) based on modified versions of previous published protocols^48–50^.

By day 20 of differentiation, hESC-derived Sinoids robustly express the SAN markers ISL1 and SHOX2:GFP, whereas ACOs and VCOs do not **Fig. 3a**). Both Sinoids and ACOs express NR2F2, while VCOs lack NR2F2 expression and instead exhibit higher levels of the ventricular-specific marker MYL2 (**Extended Data Fig. 7a**, **7b**). The differentiation efficiency is reproducible and consistent across multiple independent batches (**Extended Data Fig. 7c-7e**). Immunostaining analysis demonstrated that Sinoids contained not only SHOX2:EGFP⁺MYH6:mCherry⁺ SAN cells but also WT1⁺ epicardial cells, with CD31⁺ endothelial cells forming vessel-like structures and PHOX2B⁺ ANs located around SAN cells (**Fig. 3b, Extended Data Fig. 7f**). qRT-PCR assays confirmed the expression of representative markers of each cardiac subtype (**Extended Data Fig. 7g**). Multielectrode array (MEA) electrophysiological recordings revealed distinct membrane potential dynamics comparing Sinoids, ACOs, and VCOs over time (**Fig. 3c, Extended Data Fig. 7h, Extended Data Movies 5-7**). VCOs exhibited the slowest beating rates and more gradual changes in membrane potential, whereas Sinoids and ACOs displayed faster beating rates and more rapid membrane potential fluctuations (**Fig. 3c)**. In addition, propagation velocities differed across organoid types, with ACOs exhibiting the fastest conduction, followed by VCOs and then Sinoids (**Fig. 3d**). This hierarchy is consistent with prior knowledge that the SAN exhibits intrinsically slow conduction^51^, as well as with recent surface-based ultra–high-density electroanatomical mapping studies in juvenile pig hearts reporting higher conduction velocities in atrial working myocardium than in ventricular myocardium^52^. Functionally, pharmacological modulation of HCN4 with Ivabradine markedly reduced or abolished spontaneous electrical activity in Sinoids, as measured by MEA recordings (**Fig. 3e and Extended Data Fig. 7i**), whereas the interspike interval in ACOs remained largely unaffected (**Extended Data Fig. 7j and 7k**). These results are consistent with the SAN-specific dependence on HCN4-mediated pacemaker activity.

**Figure 3.**
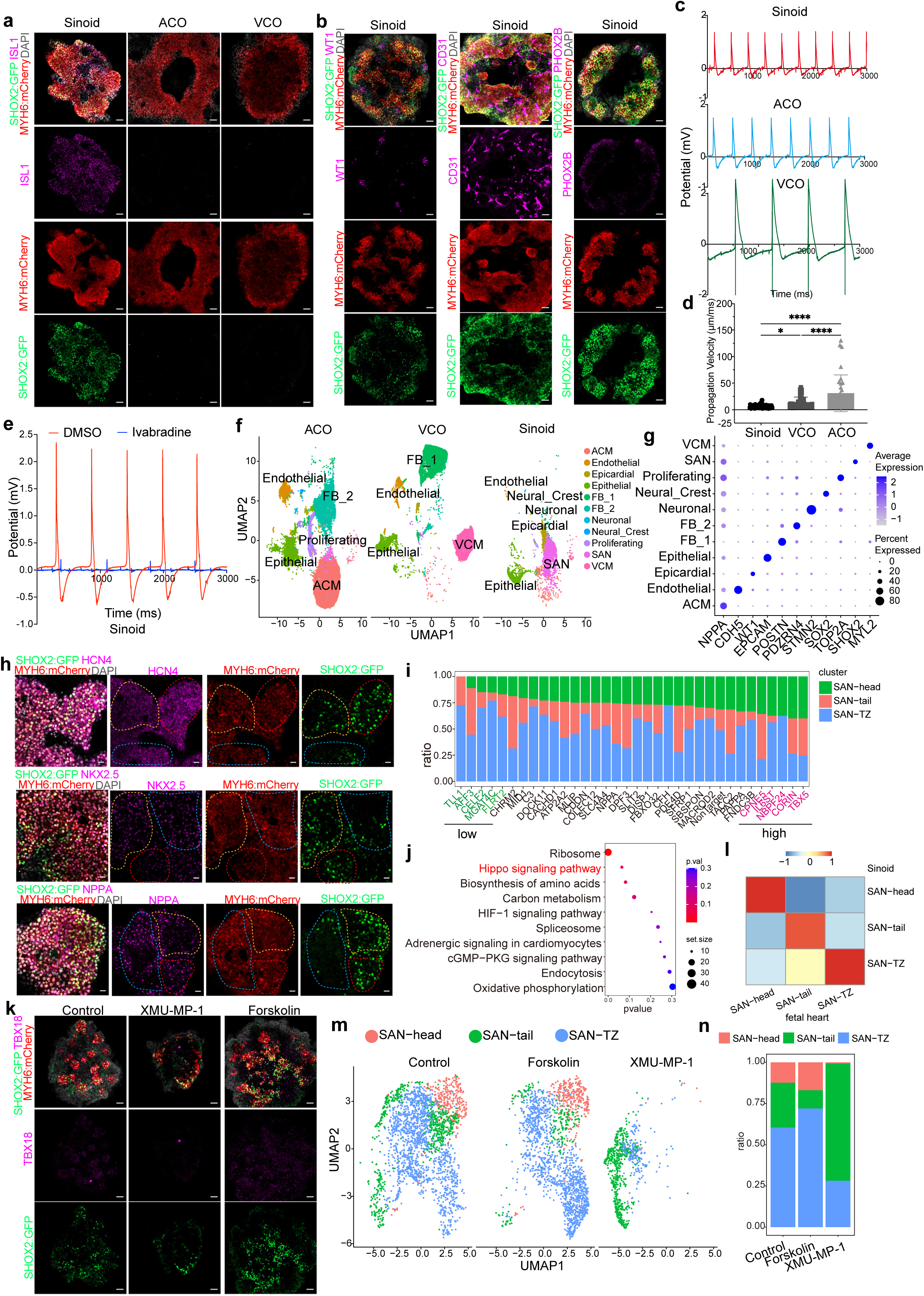
hESC-derived Sinoids recapitulate SAN function and reveal molecularly distinct SAN subpopulations. **a**, Representative confocal images of day 20 hESC-derived Sinoids, ACOs and VCOs showing expression of SAN markers. Scale bar=50 µm. **b**, Representative confocal images showing the presence of SHOX2⁺MYH6⁺ SAN cells, WT1⁺ epicardial cells, CD31⁺ endothelial cells and PHOX2B⁺ AN of day 20 hESC-derived Sinoids. Scale bar=50 µm. **c**, Representative field potential recordings from MEA showing distinct beating patterns of day 30 hESC-derived Sinoids, ACOs and VCOs. **d**, Quantification of Propagation Velocity (μm/ms) showing significant differences of Day 30 hESC-derived Sinoids, ACOs and VCOs. ****p < 0.0001; *p < 0.05. **e**, Representative MEA traces of day 30 hESC-derived Sinoids treated with 5 µM Ivabradine. **f**, UMAP plot of scRNA-seq data from day 20 hESC-derived ACOs, VCOs, and Sinoids. **g**, Dot plot illustrating expression of key marker genes across cell types within Sinoids. **h**, Representative confocal images validating SAN subcluster marker expression in hESC-derived Sinoids. **i**, Bar plots showing the distribution of SAN-head, SAN-tail, and SAN-TZ subclusters among cells carrying different sgRNAs. **j**, Pathway enrichment analysis by comparing the transcriptome changes between low-SAN-head and high-SAN-head conditions. **k**, Representative confocal images of day 20 hESC-derived Sinoids after treatment of 5 µM XMU-MP-1 or 5 µM Forskolin, Scale bar=50 µm. **l**, Correlation analysis of SAN subclusters of hESC-derived Sinoids and human fetal heart. **m**, UMAP plot of scRNA-seq data from Sinoids showing SAN subclusters. **n**, Bar plots showing the distribution of SAN-head, SAN-tail, and SAN-TZ subclusters comparing control with Forskolin or XMU-MP-1 treatment.

Next, we performed scRNA-seq to further characterize hESC-derived Sinoids, using ACOs and VCOs as reference controls. In total, we identified eleven distinct cell clusters, including *NPPA^+^* ACM, *CDH5^+^*endothelial, *WT1^+^* epicardial, *EPCAM^+^* epithelial, *POSTN^+^* fibroblast_1, *PDZRN4^+^* fibroblast_2, *STMN2^+^*neuronal, *SOX2^+^* neural crest, *TOP2A^+^* proliferating cells, *SHOX2^+^* SAN, and *MYL2^+^* VCM (**Fig. 3f, 3g, and Extended Data Fig. 8a**). Cellular composition analysis revealed that SAN cells were predominantly present in Sinoids, ACM in ACOs, VCM in VCOs, and fibroblasts abundant in both VCOs and ACOs (**Extended Data Fig. 8b-8e**). The SAN cluster exhibited high expression of markers expressed in SAN cells (*SHOX2, HCN1, HCN4, ISL1, TBX5*^53^), while the ACM cluster highly expressed atrial markers (*NPPA*, *OPCML*, *CSMD1, GJA5, ACOXL*) and the VCM cluster highly expressed ventricle markers (*HEY2*, *MYL2*, *ANKRD1, MYOZ2, IRX4*) (**Extended Data Fig. 8f**). The electrophysiological distinctions among Sinoids, ACOs and VCOs are governed by cell type–specific ion channel and gap junction expression profiles. The electrophysiological distinctions among SAN, ACMs, and ventricular VCMs are governed by cell type–specific ion channel and gap junction expression profiles. We confirmed that SAN-like cells in Sinoids express high levels of *HCN1* and *CACNA1D*, but low levels of *SCN5A*, *KCNJ2*, *GJA1* (CX43), and *GJA5* (CX40) (**Extended Data Fig. 9a**). In contrast, ACM-like cells from ACOs and VCM-like cells from VCOs exhibit high levels of *SCN5A* and *KCNJ2*. ACM-like cells are enriched for *GJA5* (CX40), while VCM-like cells predominantly express *GJA1* (CX43), supporting fast and coordinated electrical propagation in working myocardium. We also confirmed that neuronal cells expressed known AN markers (*TH*, *PRPH*, *CHAT*, *PHOX2B*, *RET*, *PHOX2A*, **Extended Data Fig. 9b**). Additional correlation analysis with published datasets of ANs, GABAergic, dopaminergic, glutamatergic, and serotonergic neurons^54,55^ confirmed that neurons in hESC-derived Sinoids most closely correlate with ANs (**Extended Data Fig. 9c**). Cell–cell interaction analysis identified conserved neuron–SAN signaling pairs, including NRG1–ERBB4, NRG3–ERBB4, and NCAM1–NCAM1, within the Sinoids (**Extended Data Fig. 9d**). These interactions closely mimic those observed in human fetal heart (**Fig. 1l**), supporting the preservation of physiologically relevant neuro–pacemaker communication in the Sinoid system.

We next evaluated the robustness of the Sinoid differentiation strategy using an independent iPSC line. This iPSC line consistently differentiated into ISL1⁺NRF2⁺ Sinoids, ISL1⁻NRF2⁺ ACOs, and *HEY2*-expressing VCOs across multiple differentiation batches (**Extended Data Fig. 10a**). qRT-PCR assays confirmed reproducible expression of lineage-specific markers across batches (**Extended Data Fig. 10b**).

### Identification of the Hippo pathway for controlling SAN subpopulation specification

Previous studies suggested that the SAN is comprised of molecularly and functionally distinct subregions, including the SAN head, SAN tail, and transitional zone (SAN-TZ)^6,56^. To assess whether these subpopulations are recapitulated in hESC-derived Sinoids, we first performed immunostaining and identified three distinct SAN cell populations: SHOX2:GFP⁺NKX2.5⁻, SHOX2:GFP⁺NKX2.5⁺, and SHOX2:GFP⁻NKX2.5⁺NPPA⁺ cells (**Fig. 3h**), corresponding to the SAN head, SAN tail, and SAN-TZ^57^, respectively. Consistent with these findings, sub-clustering of the SAN population from scRNA-seq data further confirmed the presence of three transcriptionally distinct subgroups: *SHOX2⁺NKX2.5⁻*, *SHOX2⁺NKX2.5⁺*, and *SHOX2⁻NKX2.5⁺NPPA⁺*, within hESC-derived Sinoids (**Extended Data Fig. 11a, 11b**).

UMAP visualization revealed that HCN4 expression was broadly distributed across SAN cells, whereas *SHOX2*, *NKX2.5*, and *NPPA* displayed regionally enriched expression patterns consistent with subpopulation identity (**Extended Data Fig. 11c, 11d**). Unbiased clustering analysis demonstrated that the SAN-TZ cluster is transcriptionally distinct from an ACM cluster (**Extended Data Fig. 11e**).

To identify functional regulators, we leveraged our sn-multiomics dataset from the human fetal SAN to prioritize SAN-enriched genes. We selected the top 40 SAN-enriched candidates and performed a pooled Perturb-seq screen during hESC differentiation into Sinoids. For each gene, four sgRNAs were designed (**Table S2**) and incorporated into a lentiviral CRISPR/Cas9 library. Following viral transduction and selection, cells were differentiated toward the SAN lineage and subjected to scRNA-seq analysis (**Extended Data Fig. 12a**). Analysis revealed that cells expressing sgRNAs targeting *TLL1*, *AFF3, CELF2, MGAT4C, FLRT2* are largely depleted of the SAN-head subcluster (low-SAN-head), while cells expressing sgRNAs tbx5head, **Fig. 3i**). We compared the transcriptomes changes between low-SAN-head and high-SAN-head conditions, followed by pathway enrichment analysis, which identified the Hippo signaling pathway as a top hit (**Fig. 3j**). Consistent with this analysis, dot-plot analysis confirmed high expression of Hippo pathway–related genes in cells with a high SAN-head ratio (**Extended Data Fig. 12b)**. To validate the functional role of Hippo signaling in SAN-head specification, we treated differentiating SAN cells at day 7-9 of 2D differentiation with forskolin (a chemical activator of the Hippo pathway by stimulating cAMP accumulation^58^) or XMU-MP-1 (a MST1/2 inhibitor blocking the Hippo pathway^59^) (**Extended Data Fig. 13a**). Hippo activation increased the proportions of SHOX2⁺HCN4⁺ and SHOX2⁺TBX18⁺ cells, whereas Hippo inhibition caused the opposite effect (**Extended Data Fig. 13b-13f**). With 3D organoid conditions, Sinoids derived upon XMU-MP-1 treatment exhibited a marked reduction in SHOX2⁺ cells (**Fig. 3k**). We further performed scRNA-seq to compare Sinoids generated under control, forskolin-treated, or XMU-MP-1–treated conditions. Subclustering of the SAN population identified SHOX2⁺TBX18⁺ SAN-head, SHOX2⁺NKX2.5⁺ SAN-tail, and SHOX2⁻NKX2.5⁺NPPA⁺ SAN-TZ populations (**Extended Data Fig. 14a-14c**). We also identified similar subpopulations in our spatial fetal heart sample (**Extended Data Fig. 14d**). Correlation analysis revealed a high degree of similarity between SAN subclusters in hESC-derived Sinoids and SAN cells from human fetal SAN tissue (**Fig. 3l**). Notably, the SAN-head population was almost lost in Sinoids derived upon XMU-MP-1–treated conditions, whereas forskolin treatment significantly increased the SAN-head fraction compared with controls. In addition, the SAN-TZ subcluster is also significantly depleted in XMU-MP-1–treated conditions (**Fig. 3m, 3n**). Together, these results identify Hippo signaling as a key regulatory axis controlling SAN head specification during human SAN differentiation. Several Hippo pathway associated genes also show high expression levels in SAN subcluster of human fetal SAN tissue (**Extended Data Fig. 14e**). Given that YAP is a principal downstream effector of the Hippo pathway, these findings suggest a dynamic Hippo/YAP/TEAD regulatory model, in which YAP/TEAD activity at the mesoderm stage promotes SAN differentiation, whereas activation of Hippo signaling in SAN progenitors drives specification toward the SAN head region.

### hPSC-derived SAN-paced atrial cardiomyocyte organoids (SAN-PACOs)

Spatial transcriptomics revealed that the fetal SAN directly contacts ACMs, presumably to regulate heart rhythm (**Fig. 2a**). To model this interaction *in vitro*, we assembled hESC-derived Sinoids with hESC-derived ACOs at a specific timepoint to generate SAN-paced ACOs (SAN-PACOs). After seven days of co-culture, we observed robust fusion of SHOX2-GFP⁺ Sinoids with SHOX2-GFP⁻ ACOs (**Extended Data Movies 8**). The assembly efficiency reached above 95±1.7% and was reproducible across multiple independent differentiation batches (**Extended Data Fig. 15a, 15b**). In SAN-PACOs, the Sinoid region expressed SHOX2:GFP, ISL1, and TBX18, whereas these markers were absent from the ACO region (**Fig. 4a, 4b**). CD31+ endothelial cells were present throughout the SAN-PACO (Extended Data Fig. 15c). MEA experiments were performed to assess conduction propagation in SAN-PACOs. Conduction velocity mapping revealed that electrical activity originated from the Sinoid region (**Extended Data Fig. 15d**). Treatment of SAN–PACO organoids with an HCN4 modulator abolished spontaneous electrical activity detected by MEA, reducing signals to undetectable levels (**Fig. 4c**). These results support a critical role for HCN4 in driving spontaneous electrical activity in SAN-PACOs.

**Figure 4.**
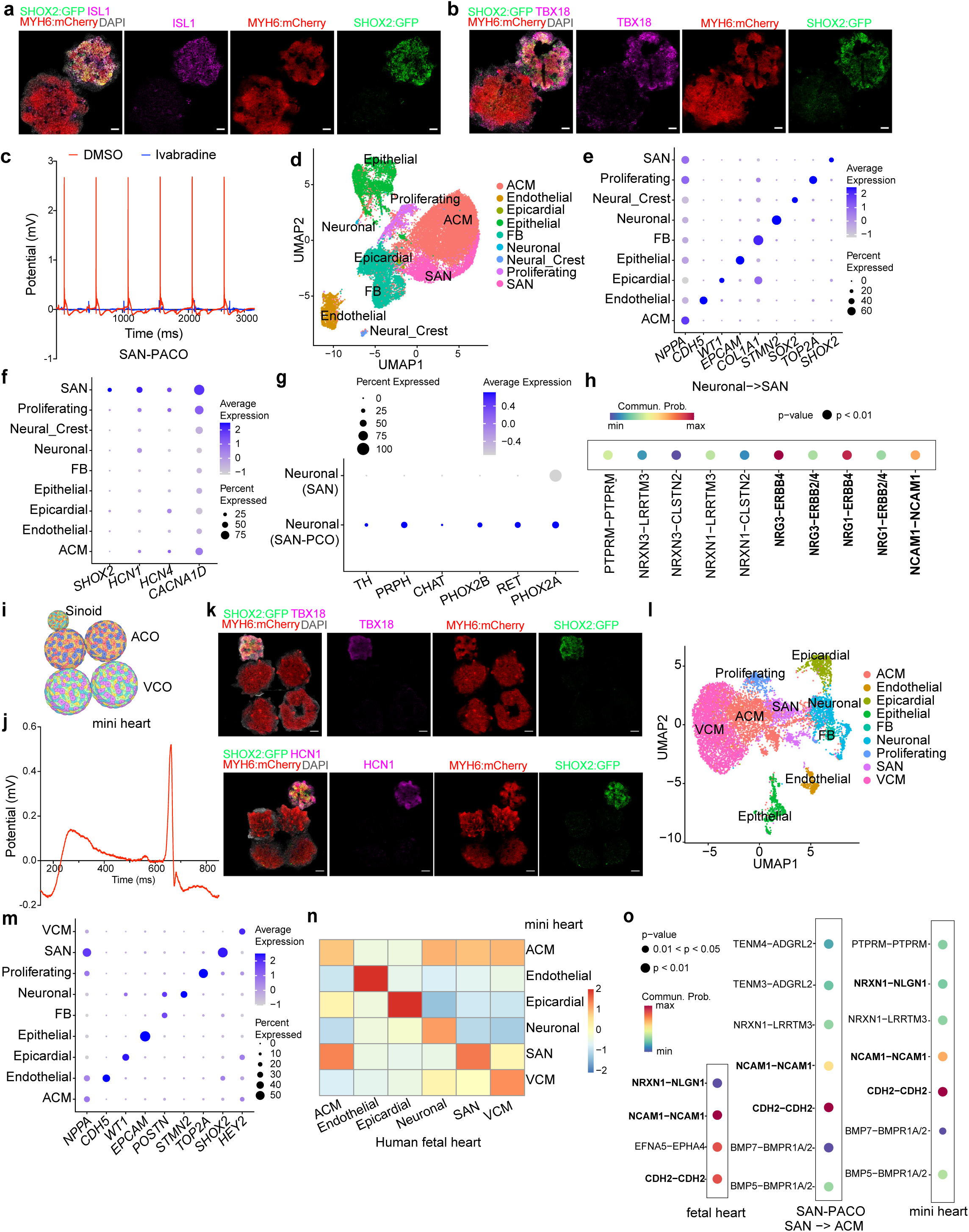
Functional and transcriptional characterization of SAN-PACOs and mini-heart organoids. **a, b,** Representative confocal images of day 20 hESC-derived SAN-PACOs showing expression of ISL1 (a) and TBX18 (b). Scale bars=50 μm. **c,** Representative MEA traces of SAN-PACOs in the presence or absence of 5 µM Ivabradine. **d,** UMAP plot of scRNA-seq data from SAN-PACOs showing distinct cell populations. **e,** Dot plot illustrating expression of key marker genes across cell types within SAN-PACOs. **f,** Dot plot illustrating expression of SAN marker genes within SAN-PACOs. **g,** Dot plot illustrating expression of AN marker genes within neuronal cluster of Sinoids and SAN-PACOs. **h,** CellChat analysis illustrating predicted neuronal-to-SAN signaling interactions. **i,** Schematic illustration showing the assembly of the mini-heart organoid. **j,** Representative field potential recordings from MEA showing distinct beating patterns of mini-heart organoids. **k,** Representative confocal images of a day 30 mini-heart organoid showing expression of ISL1 and TBX18. Scale bars=200 μm. **l,** UMAP plot of scRNA-seq data from mini heart organoids showing distinct cell populations. **m,** Dot plot illustrating expression of key marker genes across cell types within mini-heart organoids. **n,** Cross–cell-type correlation analysis comparing the human fetal heart with hESC-derived mini-heart organoids. **o,** CellChat analysis illustrating predicted SAN-to-ACM signaling interactions in human fetal heart, hESC-derived SAN-PACOs, and hESC-derived mini-heart organoids.

We performed scRNA-seq to further characterize SAN-PACOs. We identified nine distinct cell clusters, including *NPPA^+^* ACM, *CDH5^+^*endothelial, *WT1^+^* epicardial, *EPCAM^+^* epithelial, *COL1A1^+^* FB, *STMN2^+^* neuronal, SOX2*^+^*neural crest-like cells, *TOP2A^+^* proliferating cells, and *SHOX2^+^*SAN cells (**Fig. 4d, 4e, Extended Data Fig. 16a-c**). Cross-comparison with individual Sinoids and ACOs revealed that SAN cells within SAN-PACOs are predominantly contributed by Sinoids, whereas atrial ACMs in SAN-PACOs are primarily contributed by ACOs (**Extended Data Fig. 16a)**. Consistent with individual Sinoids, the SAN cluster in SAN-PACO organoids retained canonical markers expressed in SAN cells, including *SHOX2*, *HCN1*, *HCN4*, *CACNA1D*, *ISL1*, *TBX5*, *TBX18* (**Fig. 4f**) and the ACM cluster expressed atrial markers, including *NPPA*, *GJA5*, *OPCML*, *CSMD1*, *BMPER*, *ACOXL* (**Extended Data Fig. 16d**), while neuronal cells expressed established AN markers, including *TH*, *PRPH*, *CHAT*, *PHOX2B*, *RET*, *PHOX2A* (**Extended Data Fig. 16e**). Interesting, the expression levels of AN markers was significantly higher in the AN cluster of SAN-PACOs than in Sinoids alone (**Fig. 4g**) This finding is consistent with previous reports showing that cardiomyocytes secrete factors that promote AN maturation^60^. Consistent with individual Sinoids, CellChat analysis identified conserved neuron–SAN signaling pairs, including NRG1–ERBB4, NRG1–ERBB2/4, NRG3–ERBB4, NRG3–ERBB2/4, and NCAM1–NCAM1, within SAN-PACOs (**Fig. 4h**), which were highly consistent with those detected in the human fetal heart (**Fig. 1l**). Together, these findings support the preservation of physiologically relevant AN–SAN and SAN–cardiomyocyte communication within SAN-PACOs.

We also generated SAN-PACOs using an independent iPSC line, which exhibited consistent assembly efficiency across multiple batches (**Extended Data Fig. 17a, 17b**). Immunostaining further demonstrated robust enrichment of ISL1⁺ cells in the Sinoid compartment and NR2F2⁺ cells in the ACO compartment of hiPSC-derived SAN-PACOs (**Extended Data Fig. 17c**). scRNA-seq analysis confirmed that the iPSC-derived SAN-PACOs contain the major cell types, including *NPPA*^+^ ACM, *EPCAM*^+^ epithelial, *DCN*^+^ fibroblasts, *STMN2*^+^ neuronal, *TOP2A*^+^ proliferating and *SHOX2*^+^ SAN cells (**Extended Data Fig. 17d-17f**).

### hPSC-derive assembled mini-heart organoids

Building on the SAN-PACO model, we further incorporated VCOs to generate a mini-heart organoid composed of one SAN, two ACOs, and two VCOs, which together models *in vivo* heart architecture (**Fig. 4i**). The assembling efficiency of mini-hearts is around 45±2.3% across multiple batches of differentiation (**Extended Data Fig. 18a and 18b).** Following assembly and further maturation, mini-heart organoids exhibited directional beating propagation from the SAN domain to the ventricular domain (**Extended Data Movies 9**), indicative of functional pacemaker-driven conduction. Consistently, MEA recording showed a highly organized waveform with a low-amplitude pre-depolarization signal, a rapid depolarization spike, and a slower repolarization phase (**Fig. 4j**). Immunostaining demonstrated that the SAN domain of the mini-heart organoid maintains expression of SHOX2:GFP, ISL1 and HCN1, whereas these markers were absent from ACO and VCO regions (**Fig. 4k**).

To further assess cellular composition and transcriptional states, we performed scRNA-seq on mini-heart organoids. UMAP analysis identified major cardiac and non-cardiac cell clusters, including *NPPA^+^* ACM, *CDH5^+^* endothelial, *WT1^+^* epicardial, *EPCAM^+^* epithelial, *POSTN^+^*fibroblast, *STMN2^+^* neuronal, *TOP2A^+^* proliferating, *HEY2*^+^ VCM and *SHOX2^+^*SAN cells (**Fig. 4l, 4m and Extended Data Fig. 19a, 19b**). Each major cell type in the mini-heart organoid exhibited strong concordance with its counterpart in the human fetal heart based on signature gene expression, indicating faithful recapitulation of native cardiac cell identities (**Fig. 4n**). Finally, CellChat analysis revealed intercellular communication between ACM to VCM, closely resembling signaling patterns observed in the human fetal heart, thereby supporting the preservation of native cardiac signaling architecture within the mini-heart organoid (**Fig. 4o**).

### HeartOmicsAtlas web portal

To facilitate access to our spatial transcriptome, sn-multiomics and scRNA seq datasets, we developed HeartOmicsAtlas, an AI-powered web portal available at http://heartomicsdb.com. Designed for the broader research community, HeartOmicsAtlas features three main modules: Spatial Transcriptomics, sn-multiomics, and scRNA-seq. The spatial transcriptomics module allows users to visualize clustering analyses and characteristic marker expression for fetal heart, as well as explore spatially resolved gene expression across cell types. The sn-multiomics module provides clustering analyses from multiple fetal heart samples, offering tools to examine gene expression via UMAP, violin plots, and dot plots, as well as chromatin accessibility through IGV tracks. The scRNA-seq module focuses on major cell populations within hPSC-derived Sinoids, SAN-PACOs and mini-hearts, allowing users to query gene expression in annotated clusters using UMAP, violin, and dot plots. HeartOmicsAtlas provides an intuitive, AI-powered interface that supports natural language queries, enabling seamless exploration of gene expression patterns, regulatory landscapes, and cell-type composition. This interactive platform empowers researchers to investigate human heart development, dissect conduction system biology, and guide *in vitro* tissue engineering for therapeutic discovery. HeartOmicsAtlas is further enhanced by an integrated AI assistant powered by the Genomic Literature Knowledge Base (GLKB) Agent, which provides a natural language interface for intelligent data exploration. Built on a large-scale, graph-structured representation of biomedical literature and equipped with agentic reasoning capabilities, the GLKB Agent enables users to ask biologically meaningful questions and receive retrieval-grounded, literature-supported insights that connect gene expression patterns, regulatory landscapes, and cell-type composition. By combining structured multi-omics data with knowledge-graph–driven reasoning, this AI-powered interface supports both rapid exploratory analysis and deeper biological inquiry. Together, HeartOmicsAtlas and the GLKB Agent empower researchers to investigate human heart development, dissect cardiac conduction system biology, and guide in vitro tissue engineering for therapeutic discovery.

## Discussion

This study advances a conceptual and experimental framework for human SAN biology by moving beyond descriptive atlases toward functional reconstitution. Although recent single-cell analyses of fetal and adult human hearts have catalogued conduction system diversity^8–16^, including pacemaker populations, these efforts have largely remained at the level of cell-state annotation. The upstream regulatory logic that governs SAN lineage commitment, subregional specialization, and integration with working myocardium has remained unresolved. In parallel, differentiation strategies for pacemaker-like cardiomyocytes from hPSCs have relied predominantly on optimized paradigms, modulating WNT, NODAL, or retinoid signaling, or enforcing TBX3/TBX18 expression^61–68^, without systematic inference from human fetal regulatory architecture.

By integrating sn-multiomics, perturbational profiling, spatial inference, and functional tissue engineering, we define a stage-resolved regulatory architecture for human SAN development. A central insight of this work is the temporally structured role of Hippo–YAP/TEAD signaling. Early activation of YAP/TEAD promotes pacemaker lineage commitment from cardiac mesoderm, establishing SAN identity. Subsequent activation of the Hippo pathway, leading to YAP degradation, drives regional patterning toward SAN-head specification. Although YAP/TEAD signaling has previously been implicated in cardiomyocyte proliferation^69^, extracellular matrix remodeling^70^, and cardiac regeneration^71^, its role in pacemaker specification has not been previously explored in human systems. The present work therefore extends prior studies by demonstrating that YAP/TEAD signaling is not merely permissive for cardiac growth, but functions as a developmental determinant in pacemaker lineage allocation.

Beyond intrinsic transcriptional programs, we identify conserved neuron-to-SAN signaling interactions, including NRG–ERBB pathways, within the fetal SAN niche. While neuregulin signaling has been shown to influence cardiomyocyte subtype specification *in vitro*^72,73^, its integration with spatially resolved human fetal data has been lacking. By combining CellChat-based ligand–receptor inference, pharmacologic perturbation, and in vivo zebrafish validation, we demonstrate that NRG–ERBB signaling functions as a conserved regulator of pacemaker specification. The preservation of these interactions within hPSC-derived SAN organoids further supports the fidelity of the engineered system and underscores the importance of inter-lineage crosstalk in conduction system development.

Guided by these developmental insights, we established neuron-integrated SAN organoids (“Sinoids”) that recapitulate key molecular and subregional features of the fetal SAN, including SAN-head, tail, and transitional zone identities. The coordinated emergence of AN-like cells, likely neural crest–derived, within Sinoids mirrors neuron–SAN interactions observed in human fetal tissue. This co-development of mesodermal and ectodermal derivatives extends prior organoid paradigms demonstrating endoderm–mesoderm integration, including vascularized gut and lung organoids^74^ and gut organoids containing macrophages^75^. By enabling the synchronized specification of pacemaker and autonomic lineages within a single engineered platform, our system illustrates the capacity of organoid models to capture cross–germ-layer regulatory interactions that shape human cardiac conduction system development.

Assembly of Sinoids with atrial and ventricular cardioids enables the generation of SAN-PACOs and higher-order mini-hearts composed of one SAN, two ACOs, and two VCOs. Compared to previously reported multi-chamber cardioid assemblies that incorporate only atrial, left ventricular, and right ventricular compartments in a linear manner^76^, the mini-heart closely recapitulating key features of *in vivo* cardiac architecture. and a highly organized electrophysiology feature with a low-amplitude pre-depolarization signal, a rapid depolarization spike, and a slower repolarization phase. In addition, the expression of AN marker genes is higher in SAN-PACOs than in PACOs, consistent with previous studies suggesting that cardiomyocytes secrete paracrine factors that promote neuronal maturation^60^. These findings highlight the advantages of multi-organoid systems over single-organoid models in capturing inter-tissue interactions and functional maturation.

Together, this work establishes a developmental-to-engineering pipeline for human SAN biology. By defining stage-dependent regulatory determinants, reconstructing neuron–pacemaker crosstalk, and functionally integrating a pacemaker domain within multicellular cardiac assemblies, we bridge developmental genomics with human tissue engineering. These mini-heart platforms provide scalable and physiologically structured systems for modeling pacemaker dysfunction, dissecting conduction system biology, and accelerating therapeutic discovery in a human-specific context.

## Acknowledgements

This work was supported by the Department of Surgery, Weill Cornell Medicine (S.C. and T.E.).

## Author Contributions

S.C., T.E., and Y.C. conceived the study and designed the experiments. J.Z., X.D., performed multiomics and scRNA-seq library preparation, cell and organoid differentiation, FFPE tissue processing, immunostaining, qRT-PCR, and data analysis. R.G., K.B., and A.A. assisted with spatial transcriptomics library preparation, sequencing, and tissue sample processing. Z.H., K.C., K.C., K.L., and J.L. developed the HeartOmicsAtlas website and uploaded the associated code. Z.Z., performed AI-facilitated perturbation prediction. M.M., M.K., and Y.C. contributed to fetal heart sample preparation and organ collection. S.T. performed tissue sectioning and nuclear extraction for sn-multiomics and assisted with library preparation. S.C., J.Z., and T.E. wrote the manuscript.

## Methods

### Human tissue collection

All human fetal samples were collected under Institutional Review Board approval (STUDY-22-00065) at the Icahn School of Medicine at Mount Sinai. Informed consent for tissue donation was obtained by a clinical research coordinator, independent of the physician performing the procedure, only after the patient had already made the decision to terminate the pregnancy. All tissues were de-identified, and only limited clinical information was recorded, including gestational age and any maternal or fetal diagnoses.

### Visium Slide Processing and Library Preparation of Human Fetal SAN Tissue

For formalin-fixed paraffin-embedded (FFPE) fetal samples, fresh tissues were fixed in 4% (v/v) formalin at room temperature for 30 minutes, followed by three washes with 70% (v/v) ethanol. The tissues were then processed into paraffin blocks using a Tissue-Tek Vacuum Infiltration Processor 150 (Sakura Finetek) and sectioned at 10 μm thickness using a Leica microtome.

For spatial gene expression profiling, tissue sections were fixed in cold methanol and stained with hematoxylin and eosin (H&E) following 10x Genomics recommendations. Brightfield images were acquired at high resolution prior to permeabilization. Sections were permeabilized for the optimized duration to release mRNA, which hybridized to spatially barcoded oligonucleotides on the capture surface.

Following permeabilization, reverse transcription was performed in situ to generate spatially barcoded cDNA. Tissue was then removed enzymatically, and cDNA was recovered for amplification. cDNA libraries were constructed using the Visium HD Spatial Gene Expression Reagent Kit according to the manufacturer’s instructions. Library quality and fragment size distribution were assessed using an Agilent Bioanalyzer or TapeStation. Library concentration was quantified using Qubit fluorometric assays.

Prepared libraries were sequenced on an Illumina platform (NovaSeq 6000) using paired-end sequencing. Sequencing parameters followed 10x Genomics recommendations for Visium HD (28 bp for Read 1 containing spatial barcode and UMI, and 90–100 bp for Read 2 containing transcript sequence). Sequencing depth was targeted to achieve adequate coverage per capture feature as recommended by the manufacturer.

### Single nuclei extraction of human fetal SAN tissue

Flash-frozen fetal heart tissues were sectioned at 100 μm thickness and mechanically homogenized using 200 μl pipette tips in ice-cold lysis buffer (10 mM Tris-HCl pH 7.4, 10 mM NaCl, 3 mM MgCl₂, 0.1% Tween-20, 0.1% Nonidet P-40 Substitute, 1% BSA, 1 mM DTT, and 1 U/ml RNase inhibitor). The homogenate was filtered sequentially through 70 μm and 40 μm cell strainers (Corning). After centrifugation at 500 × g for 5 minutes at 4 °C, the supernatant was discarded, and the pellet was washed three times with wash buffer (10 mM Tris-HCl pH 7.4, 10 mM NaCl, 3 mM MgCl₂, 0.1% Tween-20, 1% BSA, 1 mM DTT, and 1 U/ml RNase inhibitor). Nuclei density and integrity were assessed by microscopy and were further processed for multiome paired RNA and ATAC-seq following the 10x Genomics protocol.

### Sn-multiomics library preparation of human fetal SAN tissue

Single-nucleus suspensions were adjusted to 1,000–3,000 nuclei/μl and loaded onto a Chromium Controller (10x Genomics), targeting a recovery of ∼10,000 nuclei per sample. Paired 3′ gene expression and ATAC libraries were prepared using the 10x Genomics Single Cell Multiome kit according to the manufacturer’s protocol. Library quality was assessed using the Bioanalyzer High Sensitivity DNA Analysis system (Agilent Technologies), and libraries were sequenced on an Illumina NovaSeq 6000 platform.

### scRNA library preparation of hESC-derived Sinoids, SAN-PACOs, and mini-heart organoids

Organoids were digested with Accutase (time varies, generally 30 minutes) and inverted several times every 5 minutes to obtain a single-cell suspension, washed with PBS containing 10 μM ROCK inhibitor, and filtered through a 40 μm cell strainer to remove debris and aggregates. Cell viability (>90%) and concentration (∼1,000 cells/μL) were assessed using trypan blue exclusion. Single-cell suspensions were prepared from organoids and loaded onto a Chromium Next GEM Chip using the Chromium Next GEM Single Cell 3′ HT v3.1 Kit (10x Genomics). Cells were encapsulated into GEMs for barcoding and reverse transcription. After cDNA amplification, libraries were constructed according to the manufacturer’s protocol and sequenced on an Illumina platform. Libraries were purified, quality-checked using an Agilent Bioanalyzer, and sequenced on an Illumina NovaSeq 6000 with the following read configuration: 28 bp Read 1 (cell barcode and UMI), 8 bp i7 index read, and 91 bp Read 2 (cDNA insert). Sequencing data were processed using Cell Ranger against the GRCh38 reference genome, and downstream analyses were performed in Seurat.

### Data analysis

Raw base call files were converted to FASTQ files using bcl2fastq. Sequencing reads were aligned to the human reference genome (GRCh38) using 10x Genomics Space Ranger (v4.0.1). Spatial barcode demultiplexing, UMI counting, and gene expression matrix generation were performed using default parameters. High-resolution tissue images were aligned to spatial barcode coordinates using Space Ranger image processing pipelines. Downstream analyses, including normalization, clustering, and integration with single-cell RNA-seq datasets, were performed using Seurat (5.4) and custom R scripts. Specifically, the data were normalized to account for differences in sequencing depth, followed by application of the SCTransform method, which fits regularized negative binomial models to mitigate technical artifacts while preserving biological variability. Subsequent clustering and cell-type annotation were performed following conventional scRNA-seq analysis pipelines.

#### scRNA-seq data analysis

FASTQ files were first aligned to the human genome (GRCh38) using the Cell Ranger count pipeline (v9.0.0, 10x Genomics), generating a UMI-based gene expression matrix. The resulting matrix was imported into R for downstream processing and cell cluster annotation using the Seurat package^77^. To exclude low-quality cells, potentially due to cell death or membrane damage, filtering criteria were applied based on the number of detected genes, total UMI counts, and the proportion of mitochondrial gene expression. UMI counts for each cell were normalized to the median UMI count across all cells and then log-transformed. Gene expression values were centered by subtracting the average expression of each gene across all cells. Cluster-specific positive and negative marker genes were identified by differential expression analysis relative to all other cells. Marker expression was visualized using Seurat functions including *VlnPlot, FeaturePlot,* and *DotPlot*.

#### sn-multiomics data analysis

The analysis workflow and software packages used were largely consistent with those for single-modality data. The main difference was that raw FASTQ files were processed using the Cell Ranger ARC pipeline (10x Genomics), which simultaneously generated both gene expression and chromatin accessibility (ATAC) count matrices aligned to the human genome.

#### TF enrichment analysis

To identify key transcriptional regulators, we analyzed chromatin accessibility profiles at the single-cell level using the chromVAR package. This method computes per-cell motif accessibility scores, which were incorporated into the Seurat object as a third assay (“chromvar”). Leveraging the multimodal nature of the dataset, we prioritized TFs that met independent criteria across both gene expression and chromatin accessibility analyses. Specifically, we performed two statistical tests using the Wilcoxon rank-sum method: one on gene expression and the other on chromVAR-derived motif accessibility scores. TFs with statistically significant p-values in both datasets were considered high-confidence regulators. Additionally, we calculated the area under the curve (AUC) for each gene and motif to evaluate their specificity and predictive value for individual cell types.

### 2D differentiation protocol

For differentiation, hPSCs were passaged at a density of 3 × 10⁵ cells per well in a 6-well plate and cultured for 48 hours to reach ∼80% confluence in a humidified incubator at 37°C with 5% CO₂. On Day 0, the medium was replaced with RPMI 1640 supplemented with B27 without insulin and 6 µM or 9 µM CHIR99021. On Day 2, the medium was changed to RPMI 1640 with B27 without insulin for 24 hours. On Day 3, the medium was refreshed with RPMI 1640 plus B27 without insulin containing 5 µM XAV939 plus 5 µM SB431542, 5 µM SU5402, 1 µM RA, 2.5 ng/ml BMP4, 0.1 µM Cucurbitacin I for SAN and maintained for 48 hours. On Day 5, the medium was replaced with RPMI 1640 plus B27 without insulin containing 1 µM BMS 493 for VCM or 1 µM BMS 753 for ACM or 3 µM CHIR99021, 50 ng/ml NRG1 and 10 ng/mL thyrotropin-releasing hormone (TRH) for SAN for another 48 hours. On Day 7, cells were enzymatically dissociated and replate into matrigel coated plates (Corning) to remove fibrosis cells and debris. The following day, the medium was carefully replaced with RPMI 1640 supplemented with full B27. From that point onward, the medium was refreshed every two days.

### hESC maintenance and differentiation toward Sinoids

hESC line H9, the derivative (*SHOX2:GFP; MYH6:mCherry*) line, and the iPSC line (iPSC_WCC_#8) were cultured on Matrigel-coated plates and maintained in StemFlex medium (Thermo Fisher Scientific). For differentiation, hPSCs were passaged at a density of 3 × 10⁵ cells per well in a 6-well plate and cultured for 48 hours to reach ∼80% confluence in a humidified incubator at 37°C with 5% CO₂. On Day 0, the medium was replaced with RPMI 1640 supplemented with B27 without insulin and 6 µM CHIR99021. On Day 2, the medium was changed to RPMI 1640 with B27 without insulin for 24 hours. On Day 3, the medium was refreshed with RPMI 1640 plus B27 without insulin containing 5 µM XAV939, 5 µM SB431542, 5 µM SU5402, 1 µM retinoic acid (RA), 2.5 ng/mL BMP4, 50 ng/ml NRG1, and 5 µM Kaempferol, and maintained for 48 hours. On Day 5, the medium was replaced with RPMI 1640 plus B27 without insulin containing 3 µM CHIR99021 for another 48 hours. On Day 7, cells were enzymatically dissociated and seeded into low-attachment 96-well plates (Corning) to form spheroids. The following day, the medium was carefully replaced with RPMI 1640 supplemented with full B27. From that point onward, the medium was refreshed every two days.

### hPSC differentiation toward ACOs and VCOs

For VCOs differentiation, hPSCs were passaged at a density of 3 × 10⁵ cells per well in a 6-well plate and cultured for 48 hours to reach approximately 80% confluence in a humidified incubator at 37°C with 5% CO₂. On Day 0, the medium was replaced with RPMI 1640 supplemented with B27 without insulin and 6 µM CHIR99021. On Day 2, the medium was changed to RPMI 1640 with B27 without insulin for 24 hours. On Day 3, the medium was refreshed with RPMI 1640 containing B27 without insulin, 1 µM BMS493 and 5 µM XAV939, and the cells were maintained for 48 hours. On Day 5, the medium was replaced again with RPMI 1640 supplemented with B27 without insulin plus 1 µM BMS493 and cultured for another 48 hours. On Day 7, cells were enzymatically dissociated and seeded into low-attachment 96-well plates (Corning) to form spheroids. The following day, the medium was carefully replaced with RPMI 1640 supplemented with full B27. From that point onward, the medium was refreshed every two days.

For ACOs differentiation, hPSCs were passaged at a density of 3 × 10⁵ cells per well in a 6-well plate and cultured for 48 hours to reach approximately 80% confluence in a humidified incubator at 37°C with 5% CO₂. On Day 0, the medium was replaced with RPMI 1640 supplemented with B27 without insulin and 9 µM CHIR99021. On Day 1, the medium was changed to RPMI 1640 with B27 without insulin for 48 hours. On Day 3, the medium was refreshed with RPMI 1640 containing B27 without insulin, 1 µM BMS753 and 5 µM XAV939, and the cells were maintained for 48 hours. On Day 5, the medium was replaced again with RPMI 1640 supplemented with B27 without insulin plus 1 µM BMS753 and cultured for another 48 hours. On Day 7, cells were enzymatically dissociated and seeded into low-attachment 96-well plates (Corning) to form spheroids. The following day, the medium was carefully replaced with RPMI 1640 supplemented with full B27. From that point onward, the medium was refreshed every two days.

### Generation of hPSC-derived SAN-PACO

Day 14 Sinoids and day 14 ACOs, both derived from hPSCs, were collected from low-attachment 96-well plates and combined at a 1:1 ratio. The co-cultures were maintained in RPMI 1640 medium supplemented with B27, with daily half-medium changes to minimize disturbance of the culture. SAN-PACO assembled organoids formed within 7 days, as monitored by fluorescence microscopy. Afterward, the medium was fully replaced daily to remove cellular debris and dead cells.

### Generation of hPSC-derived mini-heart organoids

Day 10 ACOs, derived from hPSCs, were collected from low-attachment 96-well plates and combined at a 1:1 ratio. Day 10 VCOs, derived from hPSCs, were collected from low-attachment 96-well plates and combined at a 1:1 ratio. Two days later, the assembled ACOs and the assembled VCOs were transferred to the same well to culture for two more days. Day 14 Sinoids were then collected and added to the previously assembled ACOs+VCOs without disturbing for two days. The final assembled mini hearts were maintained in RPMI 1640 medium supplemented with B27, with daily half-medium changes to minimize disturbance of the culture. Stable mini heart organoids formed within 10 days, as monitored by phase image. Afterward, the medium was fully replaced daily to remove cellular debris and dead cells.

### Immunofluorescence staining of cells

hPSC-derived SAN cells were fixed with 4% formalin in PBS for 30 minutes at room temperature, followed by blocking and permeabilization in 5% BSA and 0.3% Triton X-100 in PBS for 1 hour at room temperature. Cells were then incubated with primary antibodies overnight at 4 °C, followed by three washes with PBS. Secondary antibodies diluted in 5% BSA and 0.3% Triton X-100 in PBS were applied for 1 hour at room temperature. The antibody list is provided as **Table S3**. After three additional PBS washes, cell nuclei were stained with DAPI for 20 mins at room temperature. Finally, cells were rinsed three times in PBS before imaging with a Zeiss LSM-780 inverted confocal microscope.

### Immunofluorescence staining of Sinoids, SAN-PACO, and mini-heart organoids

hPSC-derived Sinoids, SAN-PACOs and mini-heart organoids were collected into Eppendorf tubes, and the supernatant was carefully removed. Samples were fixed in 4% formalin in PBS for 30 mins at room temperature, followed by blocking and permeabilization in 5% BSA and 0.3% Triton X-100 in PBS overnight at 4 °C. Organoids were incubated with primary antibodies for 48 hours at 4 °C, followed by three washes with PBS. Secondary antibodies, diluted in 5% BSA and 0.3% Triton X-100 in PBS, were applied for 48 hours at 4 °C. After three additional PBS washes, cell nuclei were stained with DAPI for 30 minutes at room temperature. Organoids were rinsed three times with PBS and transferred to confocal imaging dishes for image acquisition.

### Perturb-seq analysis

To perform Perturb-seq, we first incorporated random 22-nucleotide genetic barcodes (GBCs) into the backbone plasmid pPS. The pooled sgRNA mixture was PCR amplified and ligated into this plasmid. The sequences of sgRNAs are provided in **Table S2**. The sgRNA-GBC correlation reference table was generated via sequencing. An EGFP coding sequence driven by the EF1α promoter was then inserted between the sgRNA and GBC elements. The final construct was packaged into lentivirus using 293T cells. Separately, a plasmid encoding CRISPR/Cas9 and co-expressing mCherry was also packaged into lentivirus.

hPSCs were co-infected with both lentiviral constructs and sorted based on fluorescent protein expression. After a one-week recovery period, the cells were induced to differentiate toward the SAN lineage. On day 20 of differentiation, SAN cells were harvested and prepared for scRNA-seq according to the 10x Genomics Single Cell Protocols Cell Preparation Guide. Cells were encapsulated into droplet emulsions using the Chromium Controller and Chromium Single Cell 3’ Gel Beads v3 (10x Genomics), aiming to recover ∼10,000 cells per GEM group.

To establish the correlation between cell barcodes (CBCs) and GBCs, an additional round of enrichment PCR and sequencing was performed using intermediate products from the library preparation. For perturbation analysis, each cell was annotated with its corresponding perturbation target based on the sgRNA-GBC and CBC-GBC reference tables. Cells harboring multiple perturbations were excluded from analysis. Finally, we assessed the impact of each gene perturbation by examining shifts in the distribution of SAN subpopulations. Pathway enrichment analysis was performed using the GAGE R package^78^. Normalized gene expression data were analyzed against curated gene sets from KEGG/MSigDB. GAGE was used to identify significantly upregulated or downregulated pathways between experimental conditions. P-values were adjusted using the Benjamini–Hochberg method, and pathways with FDR < 0.05 were considered significant.

### MEA analyses

Spontaneous field potentials (FPs) were recorded from Sinoids, ACOs, VOCs, SAN-PACO, and mini-heart organoids using the MED64 MEA system (AlphaMED Science, Osaka, Japan). Organoids were transferred to MEA probes, and FPs were measured immediately. For drug treatment experiments, the culture medium within the MEA probes was replaced with drug-containing medium, and recordings were initiated immediately after the medium change. All measurements were conducted at 37 °C in a humidified environment with 7.5% CO₂ using a TOKAI HIT incubator (Shizuoka, Japan).

### qRT-PCR assays

Total RNA was extracted and purified using the RNeasy Plus Kit (Qiagen), followed by reverse transcription using the High-Capacity cDNA Reverse Transcription Kit supplemented with RNase inhibitor (Thermo Fisher). Quantitative PCR was performed using SYBR Green Gene Expression 2X Mix (Vazyme) and gene-specific primers. Expression levels were normalized to the human housekeeping gene *TBP* as an internal reference. Primer sequences are provided in **Table S4**.

### Zebrafish pacemaker analysis

The Tg(*nkx2.5:ZsYellow*)^fb7^ reporter strain^79^ was kindly provided by the Burns laboratory (Children’s Hospital, Boston, MA). All animal work was performed under an approved IACUC protocol from Weill Cornell Medicine with animal care under supervision of the Research Animal Resource Center. Larvae were obtained by natural matings of adult transgenic animals and ZsYellow+ embryos identified manually at 24 hpf. Starting at 30 hpf, manually dechorionated larvae were placed in petri dishes with E3 buffer containing 1% DMSO, 1 mM Tris ph7.5 and exposed to 5 µM Afatinib or BAY-593 (from 10 mM stocks in DMSO), or an equivalent volume of DMSO as control. At 54 hpf larvae were fixed in 2% PFA for 2 hours at room temperature, washed extensively in PBS with 0.3% TritonX-100 (PBX), incubated in blocking buffer (BB: PBX, 0.5% BSA, 10% goat serum) for 2 hours at room temperature, incubated overnight at 4°C in BB containing 1/50 dilution of mouse anti-ISL1/2 antibody (Iowa Developmental Studies Hybridoma Bank #39.4D5), washed extensively with PBX, incubated at room temperature for 2 hours in BB with a 1/200 dilution of goat anti-mouse IgG2b Alexa Fluor568 (Invitrogen #A-21144), washed extensively with PBX, and mounted in 1:1 PBS:glycerol for imaging on a Zeiss LSM800 confocal microscope. Individual Isl1+ cells were counted by a blinded investigator from imaging through Z-stacks. The areas of Isl1+ regions were quantified using ImageJ.

### HeartOmicsAtlas

HeartOmicsAtlas features an advanced AI assistant designed to improve accessibility and streamline user interaction through a dual-component, LLM-based architecture. The first component is a data interaction agent, powered by the ChatGPT-4o model, which enables natural language queries, demystifies complex biological terminology, facilitates access to experimental datasets, and provides concrete examples of platform functionality. This agent focuses on direct interaction with HeartOmicsAtlas, interpreting user intent through carefully designed prompts and dynamically invoking backend APIs to retrieve and visualize relevant spatial transcriptomics, sn-multiomics, and scRNA-seq data. By tightly integrating large language model capabilities with the platform’s API infrastructure, this component delivers accurate, context-aware, and responsive support for interactive data exploration, while remaining optimized for scalability and concurrent user queries.

Complementing this data-centric agent, HeartOmicsAtlas integrates a GLKB Agent as a second, specialized AI component that provides literature-grounded biological reasoning and contextual interpretation. The GLKB Agent^80^ is built on the Genomic Literature Knowledge Base (GLKB), a large-scale, graph-structured representation of biomedical literature, and is designed to support complex biological queries that extend beyond direct dataset visualization. When user questions require mechanistic interpretation, cross-study evidence synthesis, or broader biological context, such as gene regulatory relationships, disease relevance, or developmental pathways, the GLKB Agent performs structured retrieval and multi-hop reasoning over curated literature to generate evidence-supported responses. Importantly, the GLKB Agent does not replace the data interaction agent; instead, it operates as a complementary reasoning module, enriching HeartOmicsAtlas with literature-based insights while preserving a clear separation between data access and visualization and knowledge-driven biological interpretation.

### Quantification and Statistical analysis

N=3 independent biological replicates were used for all experiments unless otherwise indicated. n.s. indicates a non-significant difference. P-values were calculated by one-way analysis of variance (ANOVA) unless otherwise indicated. *p<0.05, **p<0.01, ***p<0.001 and ****p<0.0001.

## Data availability

Sn-multiomics and scRNA-seq has been uploaded to GEO. GSE301201, GSE301202, GSE301203.

## Code availability

The analysis was performed using standard protocols with previously described computational tools. The codes are uploaded to the GitHub: https://github.com/JerryJiajun/fetal_heart.git. https://github.com/HowIII/fetal_heart_insilico_perturbation https://github.com/Kevin4335/HeartOmicsAtlas-public.

## Extended Data Movies

**Movie 1: Live imaging of hESC-derived cells control.**

**Movie 2: Live imaging of hESC-derived cells overexpressing *MEF2A*.**

**Movie 3: Live imaging of hESC-derived cells overexpressing *MGA*.**

**Movie 4: Live imaging of hESC-derived cells overexpressing *TEAD1*.**

**Movie 5: Live imaging of hESC-derived ACO.**

**Movie 6: Live imaging of hESC-derived VCO.**

**Movie 7: Live imaging of hESC-derived Sinoid.**

**Movie 8: Live imaging of hESC-derived SAN-PACO.**

**Movie 9: Live imaging of hESC-derived mini-heart organoid.**

---

**Extended Data Figure 1.**
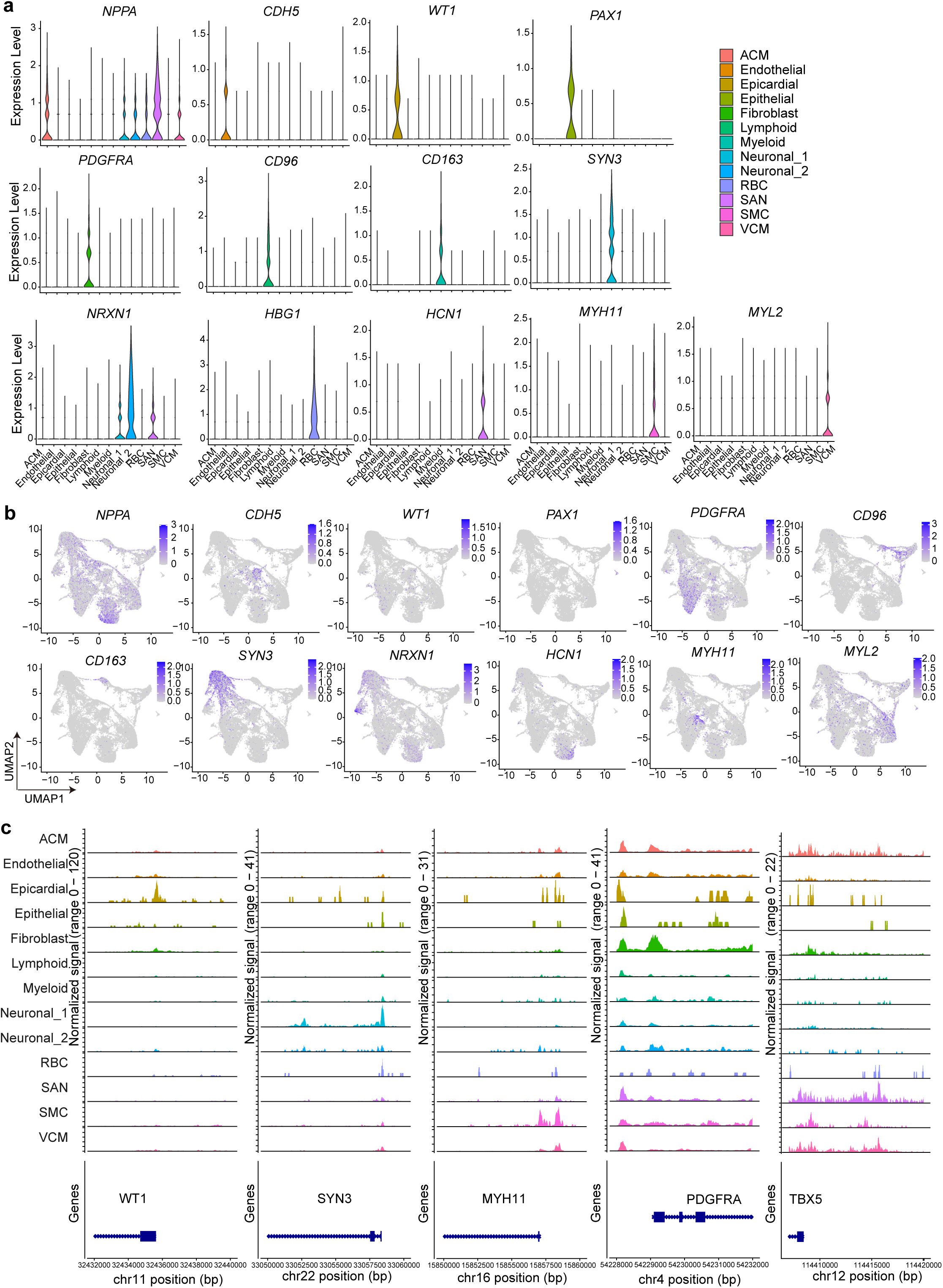
Characterization of cell types in the human fetal SAN by sn-multiomics profiling. **a,** Violin plots showing the expression distribution of representative cell type–specific markers. **b,** UMAP plots displaying the expression patterns of selected marker genes. **c,** IGV tracks illustrating chromatin accessibility at loci near representative cell type–specific genes.

**Extended Data Figure 2.**
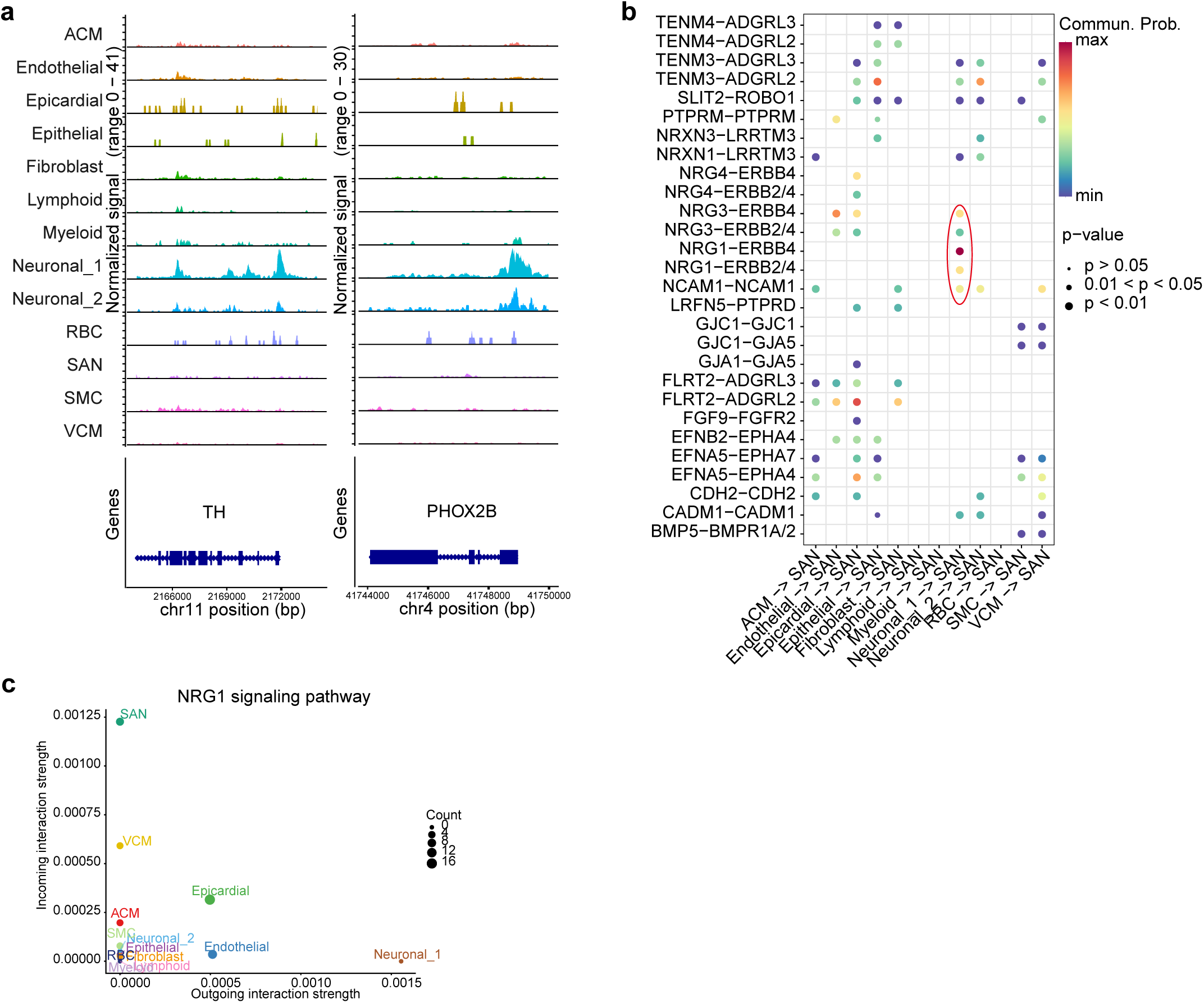
Cell–cell interactions in human fetal SAN tissue. **a,** IGV tracks illustrating chromatin accessibility at loci near representative AN genes. **b,** Bubble plot showing significant signaling interactions from other cell populations to SAN cells. **c,** Signaling role analysis of the aggregated cell–cell communication network for the NRG signaling pathway.

**Extended Data Figure 3.**
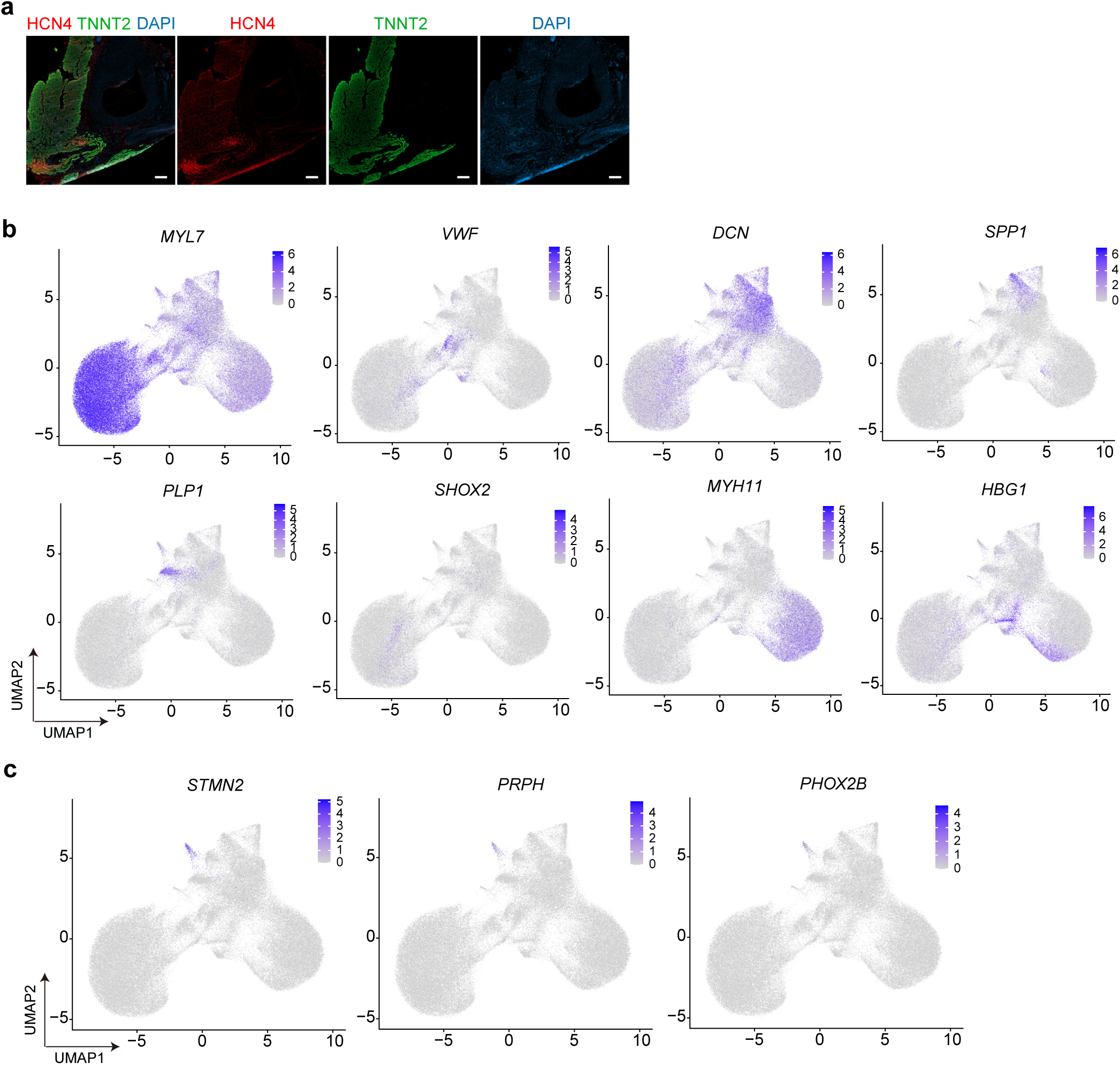
Characterization of cell types in the human fetal SAN by spatial transcriptome profiling. **a,** Confocal IF images showing the expression of HCN4 and TNNT2 on the tissue slide. Scale bar=100 µm. **b,** UMAP plots illustrating the expression patterns of select marker genes. **c,** UMAP plots illustrating the expression patterns of AN marker genes.

**Extended Data Figure 4.**
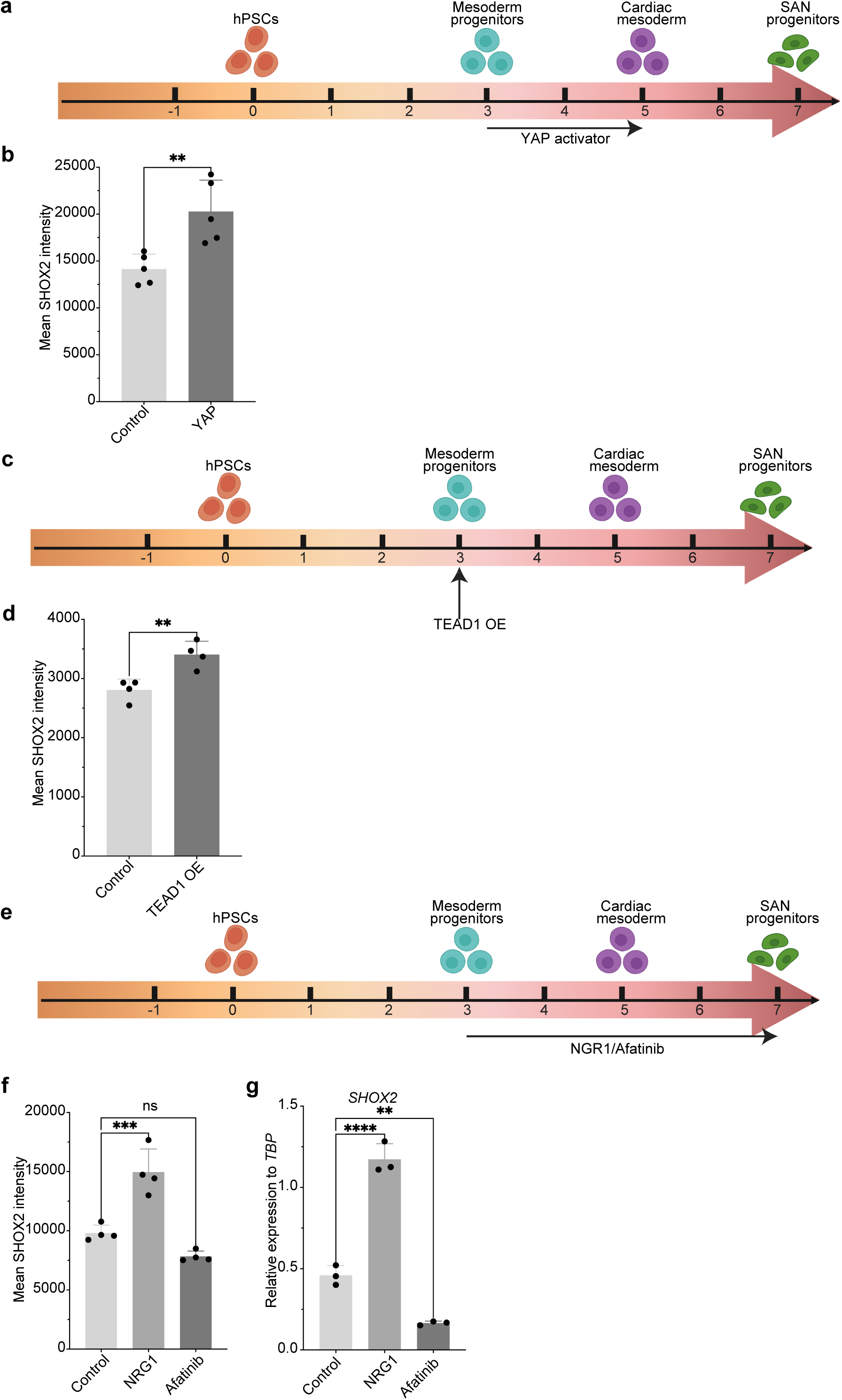
Characterization of hESC-derived SAN after modifying YAP/TEAD or NRG pathways. **a,** Schematic of kaempferol treatment during 2D hESC SAN differentiation. **b,** Quantification of mean SHOX2 intensity of day 20 hPSC-derived SAN after 5 µM kaempferol or DMSO control treatment during differentiation. Data are shown as mean ± SEM; **p < 0.01; N = 3 biological replicates. **c,** Schematic of TEAD1 overexpression (OE) during 2D hESC SAN differentiation. **d,** Quantification of mean SHOX2 intensity of day 20 hESC-derived SAN cells with control or *TEAD1* OE. Data are shown as mean ± SEM; **p < 0.01; N = 3 biological replicates. **e,** Schematic of NRG1 or Afatinib treatment during 2D hESC SAN differentiation. **f,** Quantification of mean SHOX2 intensity of day 20 hESC-derived SAN cells following 50 ng/ml NRG1 or 5 µM Afatinib treatment compared to DMSO control. Data are shown as mean ± SEM; ***p < 0.001; N = 3 biological replicates. **g,** qRT-PCR of *SHOX2* expression of day 20 hESC-derived SAN cells following 50 ng/ml NRG1 or 5 µM Afatinib treatment compared to DMSO control. Data are shown as mean ± SEM; **p < 0.01; ****p < 0.0001; N = 3 biological replicates.

**Extended Data Figure 5.**
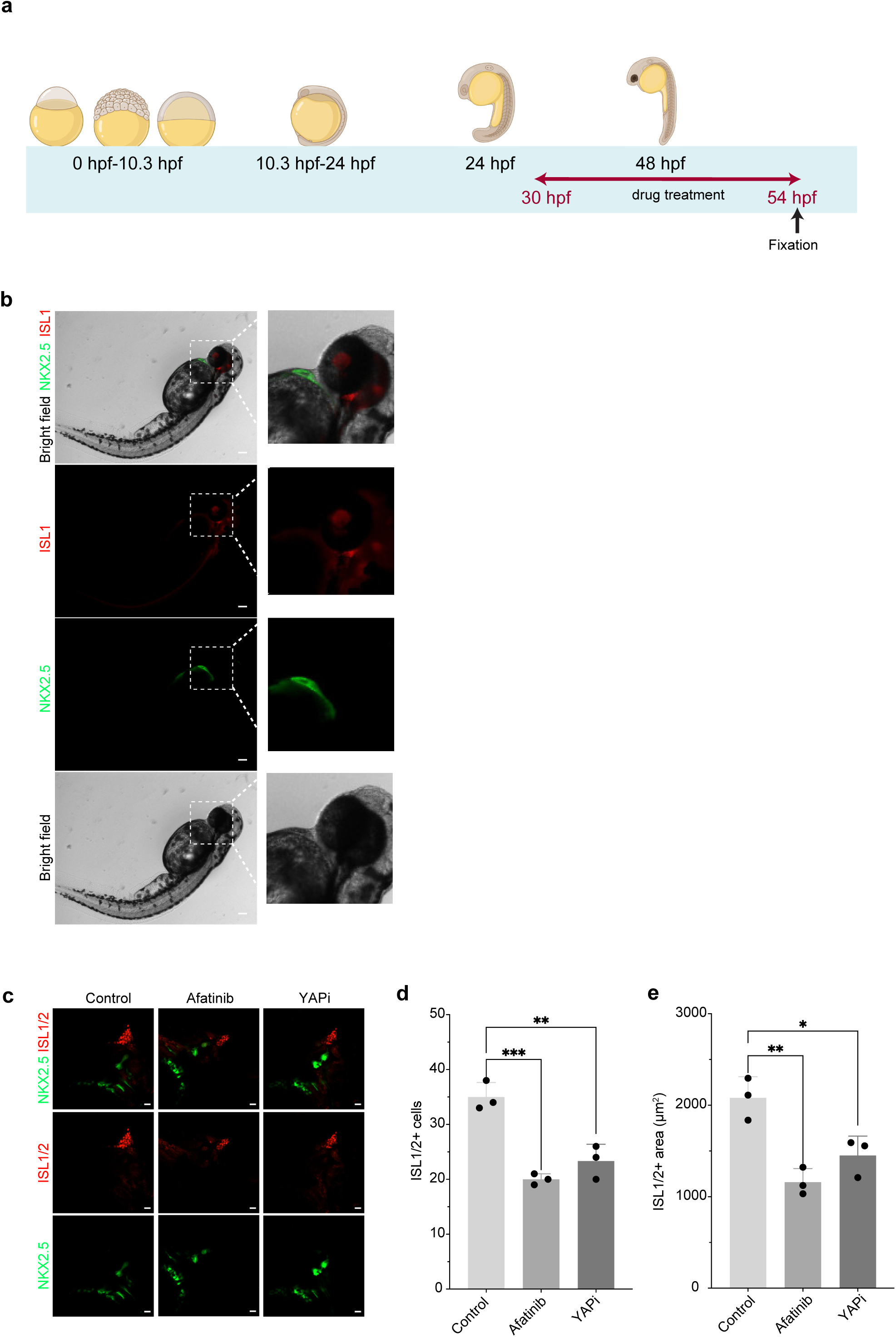
*In vivo* analysis of SAN development in zebrafish following drug treatments. **a,** Schematic of zebrafish drug treatment experiments. **b,** Representative bright-field images of zebrafish larvae at 54 hpf. Scale bar=100 µm. **c,** Representative fluorescence images of zebrafish /larvae at 54 hpf treated with 5 µM Afatinib, 5 uM BAY-593 (YAPi) or vehicle control. Scale bar=100 µm. **d, e,** Quantification of the number of ISL1/2^+^ cells (c) and ISL1/2^+^ area in zebrafish larvae treated with 5 uM Afatinib, 5 uM BAY-593 or vehicle control. Data are shown as mean ± SEM; *p < 0.05; **p < 0.01; ***p < 0.001; ****p < 0.0001. For each experiment, N = 25 larvae. For each drug-treated set there was a consistent loss of Isl1-staining; 3 larvae from each set were randomly chosen for quantification, carried out by a blinded investigator.

**Extended Data Figure 6.**
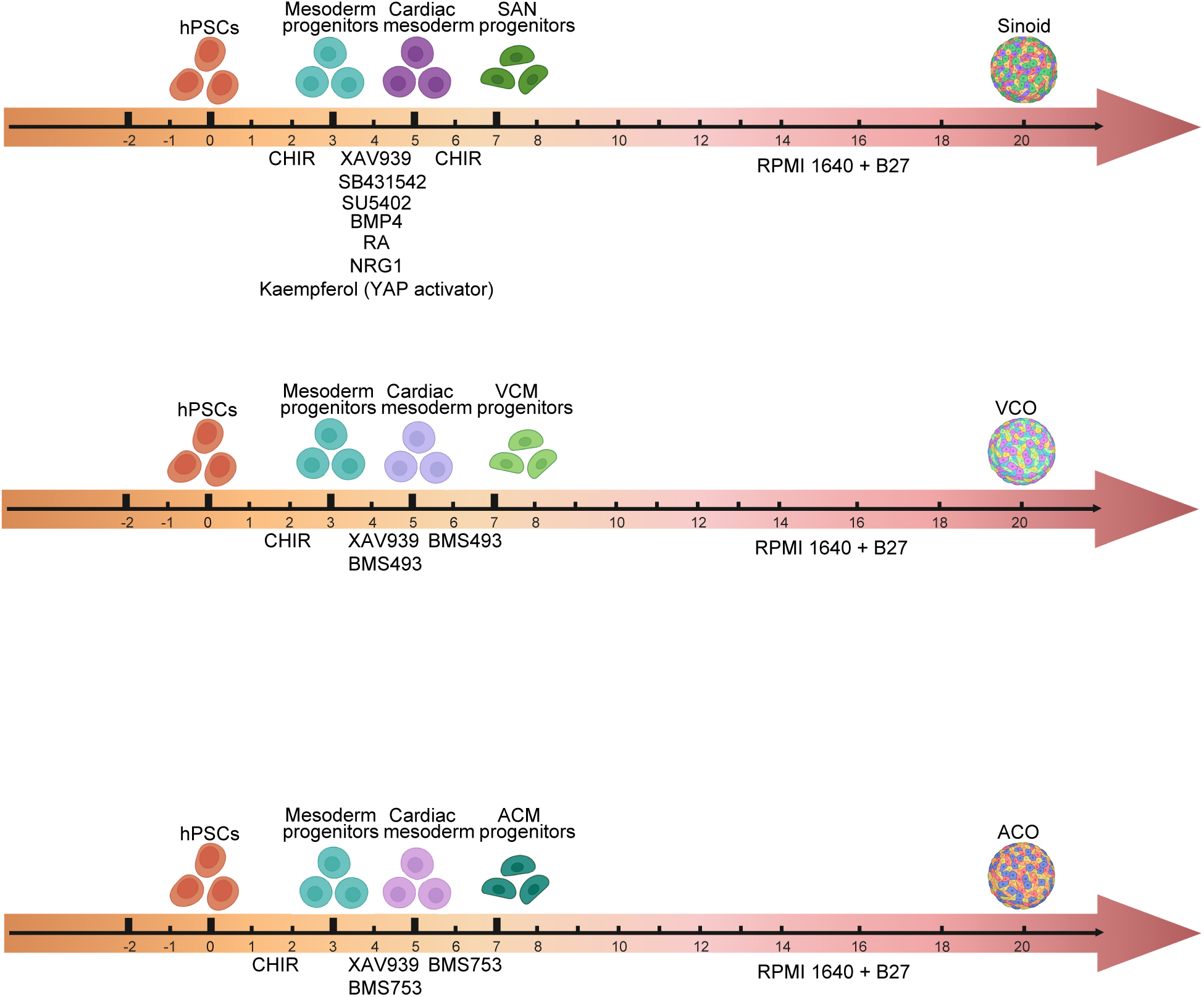
**Schematic of the stepwise protocol used to differentiate hPSCs into 3D Sinoids, ACOs, or VCOs.**

**Extended Data Figure 7.**
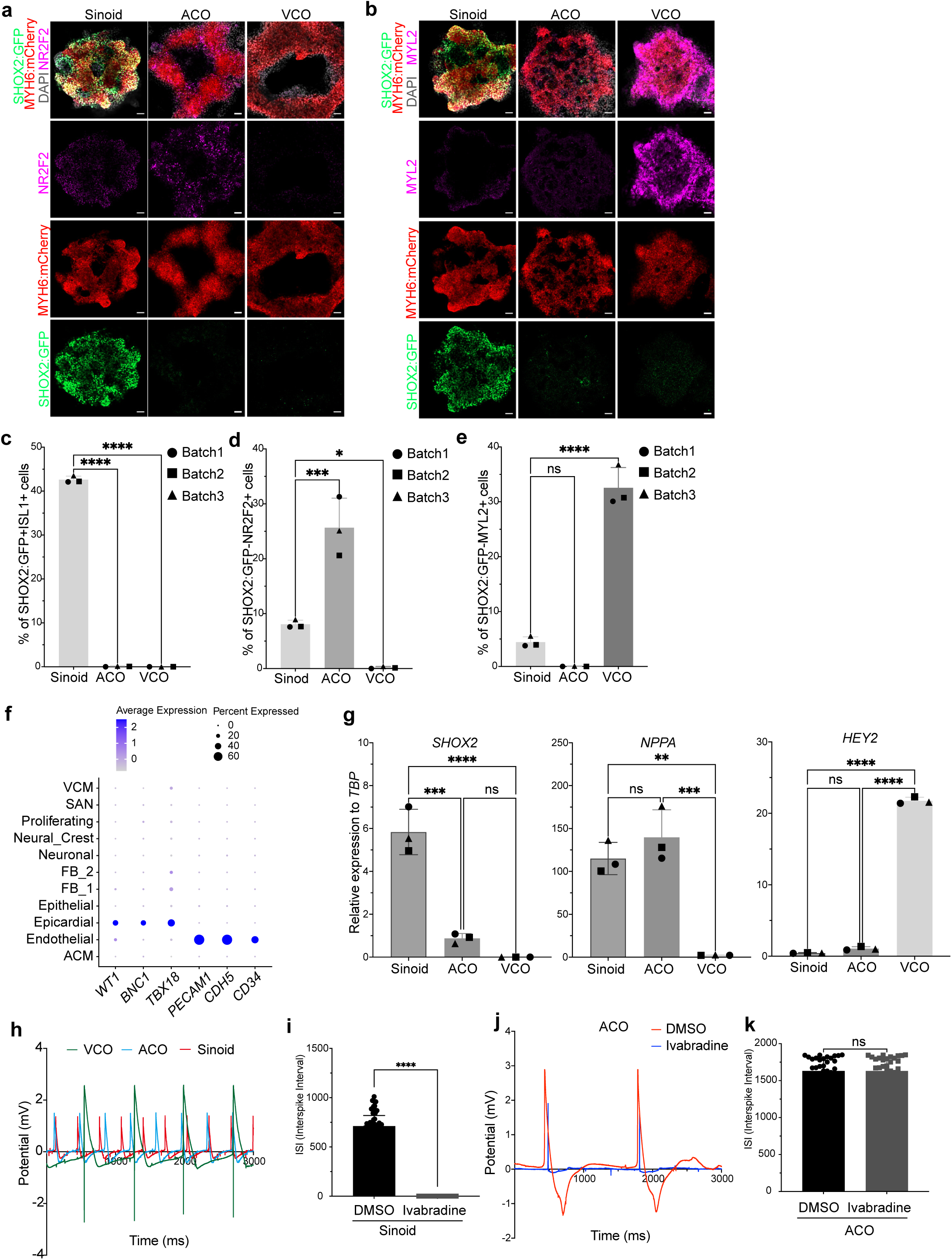
Characterization of hESC-derived Sinoid, ACOs and VCOs. **a, b,** Representative confocal images of day 20 hESC-derived Sinoids, ACOs, and VCOs stained for NR2F2 (a) or MYL2 (b). Scale bar=50 µm. **c-e,** Quantification of the percentage of SHOX2:GFP^+^ISL1^+^ cells (c), SHOX2:GFP^-^NR2F2^+^ cells (d) and SHOX2:GFP^-^ MYL2^+^ in day 20 hESC-derived Sinoids, ACOs, and VCOs across multiple independent differentiation batches. *p < 0.05; ***p < 0.001; ****p < 0.0001. **f,** Dot plot illustrating the expression of epicardial and endothelial cell markers in hESC-derived Sinoids. **g,** qRT-PCR analysis of day 20 hESC-derived Sinoids, ACOs, and VCOs across multiple independent differentiation batches. *p < 0.05; ***p < 0.001; ****p < 0.0001. **h,** Representative MEA traces of day 20 hESC-derived Sinoids, ACOs, and VCOs. **i,** Quantification of interspike interval of day 20 hESC-derived Sinoids treated with vehicle control or 5 µM Ivabradine. **j,** Representative MEA traces of hESC-derived ACOs treated with vehicle control or 5 µM Ivabradine. **k,** Quantification of interspike interval of day 20 hESC-derived ACOs treated with vehicle control or 5 µM Ivabradine.

**Extended Data Figure 8.**
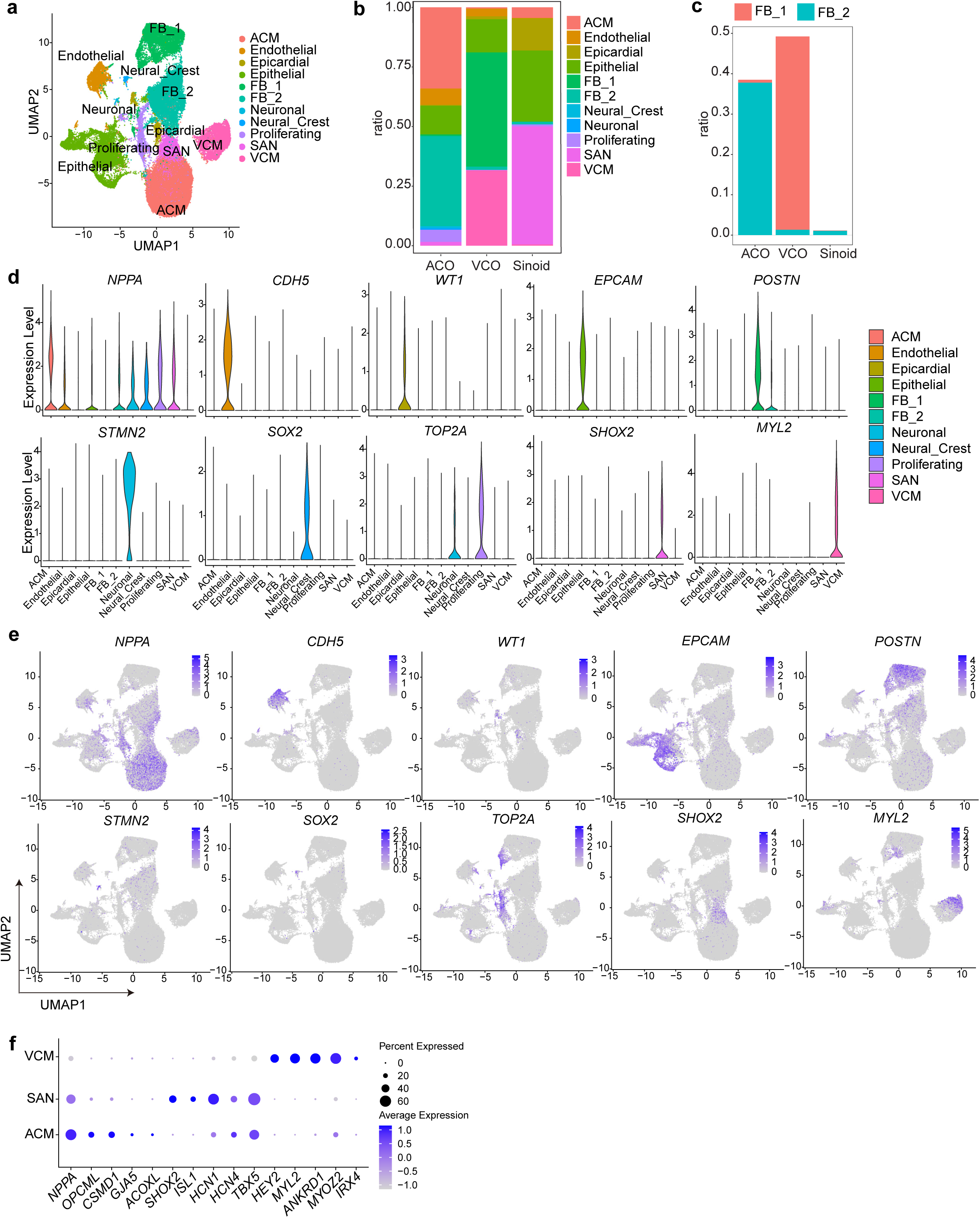
scRNA-seq of hESC-derived Sinoids, ACOs, and VCOs by scRNA-seq. **a,** Integrative UMAP of hESC-derived Sinoids, ACOs, and VCOs. **b,** Bar plot displaying the relative proportions of each cell type in hESC-derived Sinoids, ACOs, and VCOs. **c,** Percentage of fibroblast cells in hESC-derived Sinoids, ACOs, and VCOs. **d,** Violin plots illustrating the expression levels of representative cell type–specific marker genes. **e,** UMAP plots illustrating the expression patterns of cell type–specific markers. **f,** Dot plot illustrating the expression of atrial, ventricular, and SAN marker genes across ACM from ACO, VCM from VCO, and SAN from Sinoid populations.

**Extended Data Figure 9.**
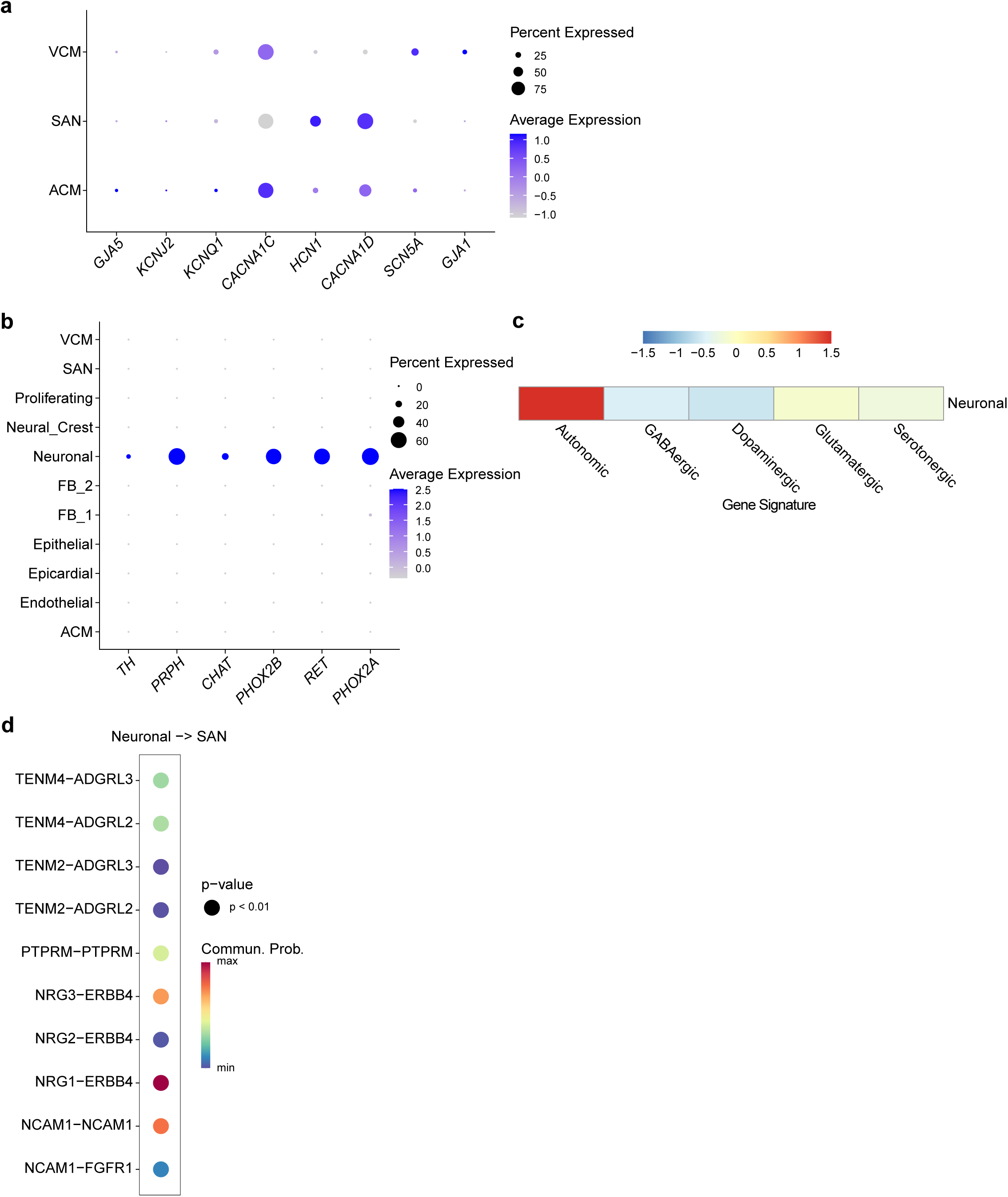
CellChat analysis of hESC-derived Sinoids. **a,** Dot plot illustrating the expression of genes encoding ion channels within SAN-like, ACM-like and VCM-like clusters of hESC-derived Sinoids. **b,** Dot plot illustrating the expression of AN genes within neuronal cluster of hESC-derived Sinoids. **c,** Correlation analysis between neuronal cluster of hESC-derived Sinoids with published datasets of ANs, GABAergic, dopaminergic, glutamatergic, and serotonergic neurons. **d,** CellChat analysis illustrating predicted Neuronal-to-SAN signaling interactions.

**Extended Data Figure 10.**
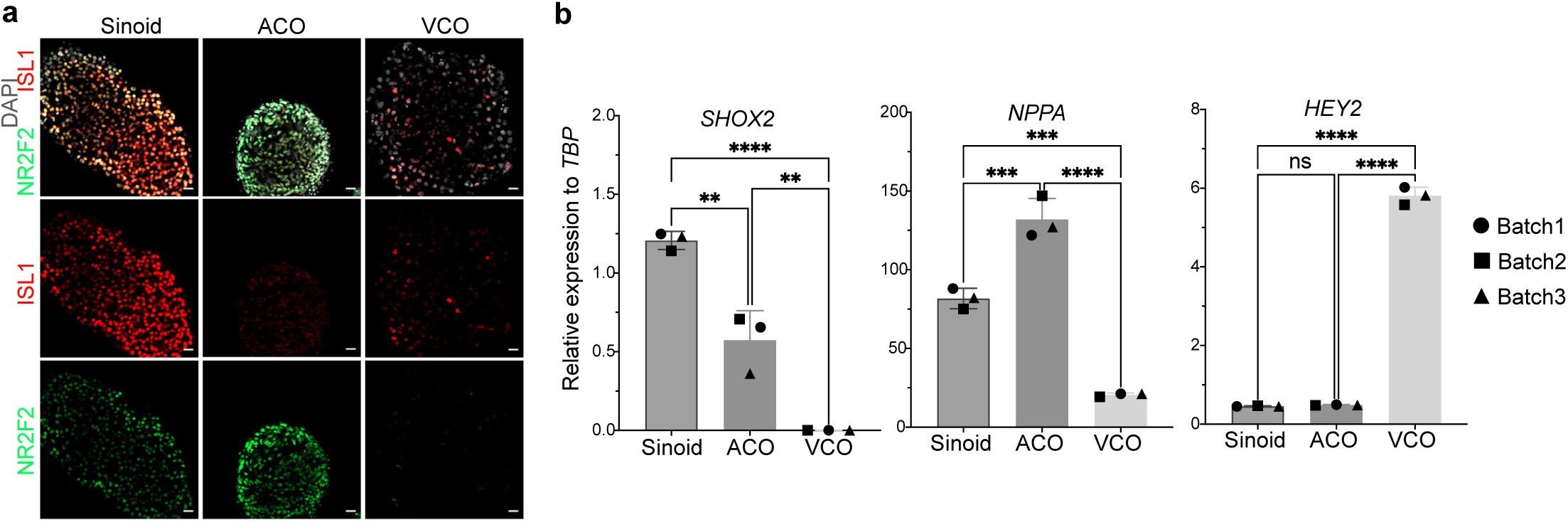
Characterization of hiPSC-derived sinoids. **a**, Representative confocal images of day 20 hiPSC-derived Sinoids, ACOs and VCOs showing expression of SAN markers. Scale bar=50 µm. **b,** qRT-PCR analysis of marker genes expression in day 20 hiPSC-derived Sinoids, ACOs, and VCOs across multiple independent differentiation batches. Data are shown as mean ± SEM; ****p < 0.0001; ***p < 0.001; **p < 0.01.

**Extended Data Figure 11.**
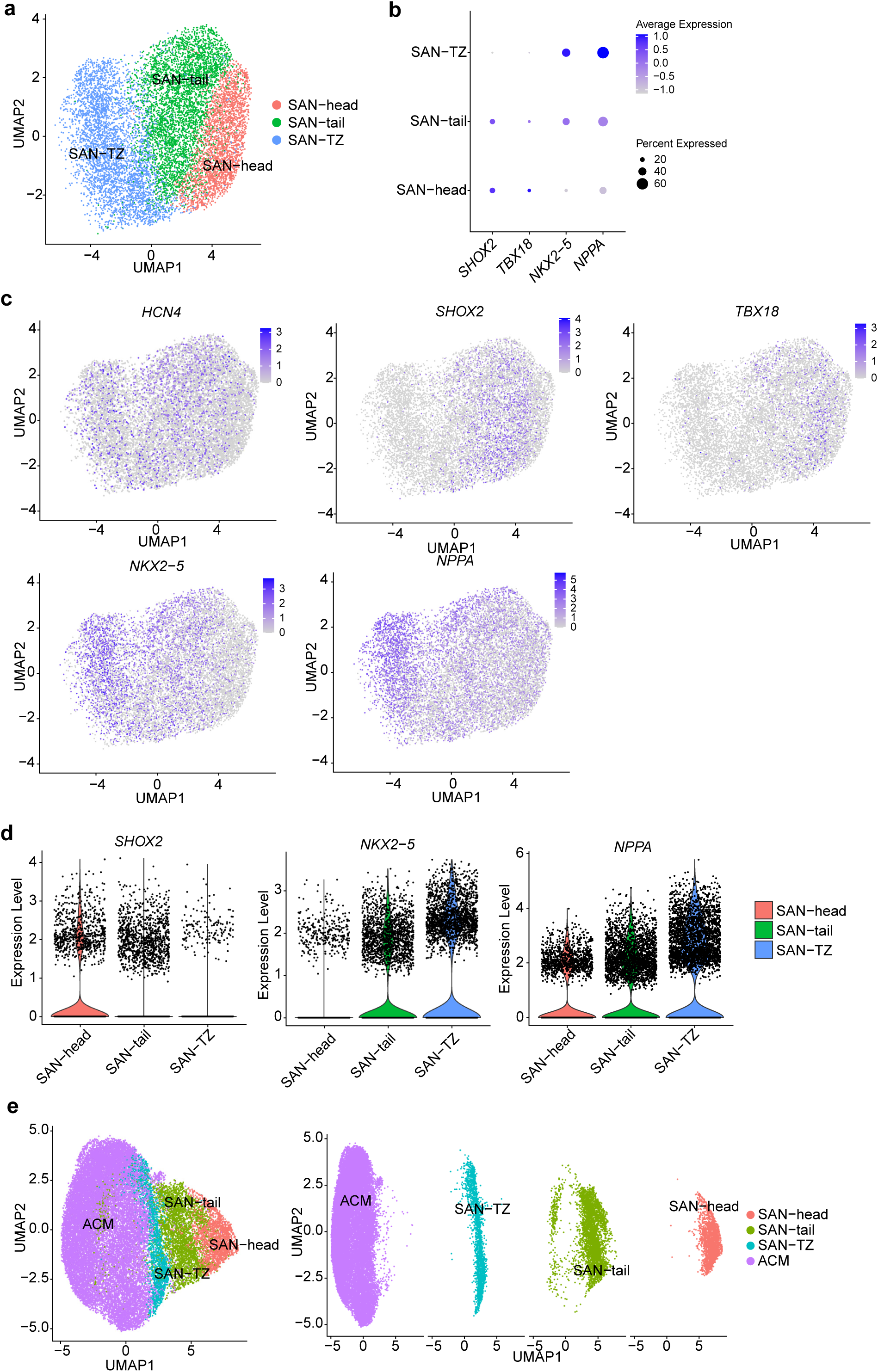
Characterization of subclusters within the SAN population of hESC-derived Sinoids. **a,** UMAP plot displaying composition of subclusters within SAN population. **b,** Dot plots showing expression levels of subcluster–specific markers. **c,** UMAP plot displaying expression of subcluster–specific markers. **d,** Violin plots showing expression levels of subcluster–specific markers. **e,** UMAP showing distinguished cluster of ACM coming from ACO, SAN-TZ, SAN-tail, and SAN-head coming Sinoid.

**Extended Data Figure 12.**
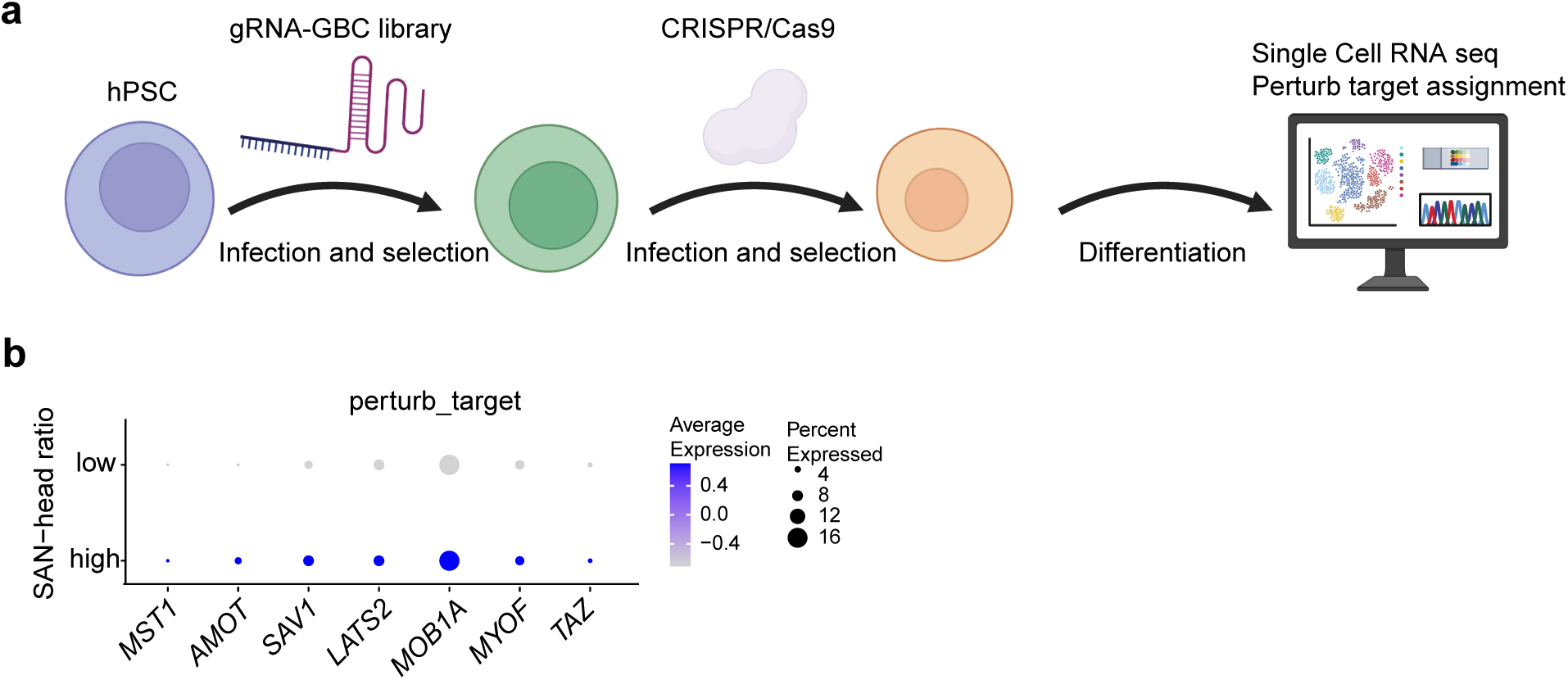
Perturb-seq. **a,** Schematic overview of the Perturb-seq workflow. **b,** Dot plot showing expression of representative Hippo pathway related genes.

**Extended Data Figure 13.**
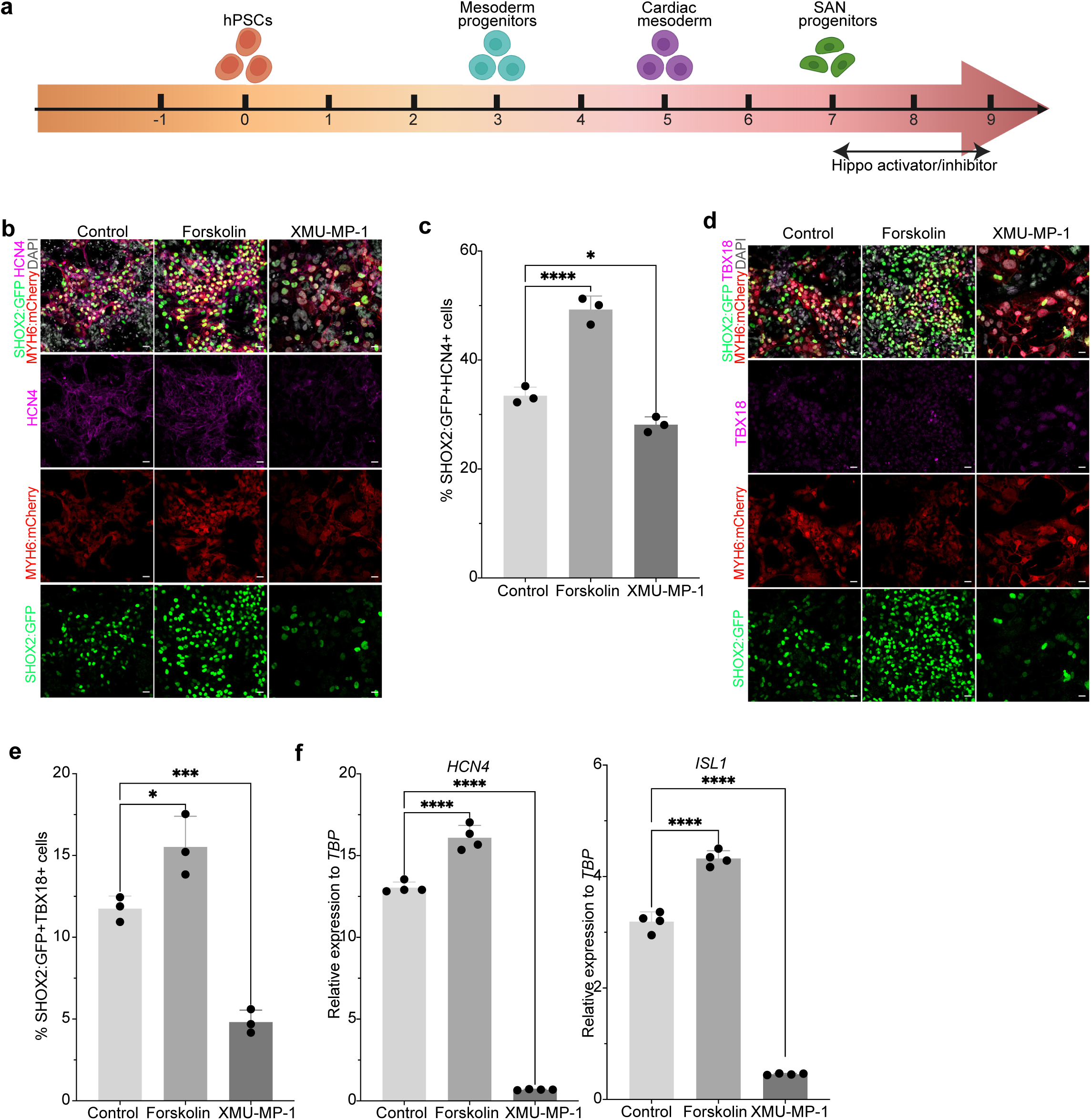
Hippo pathway modulation regulates SAN-head specification in hESC-derived Sinoids. **a,** Scheme of the treatment with 5 µM Forskolin or 5 µM XMU-MP1 (day 7 to day 9) to evaluate impact on SAN subcluster specification. **b**, Representative confocal images showing expression of SHOX2, HCN4, and MYH6 following treatment with the indicated drugs. Scale bar=20 µm. **c**, Quantification of the percentage of SHOX2^+^/HCN4^+^ cells in control, forskolin and XMU-MP-1 treated cells, respectively. ****p < 0.0001; *p < 0.05. Data are shown as mean ± SEM. N = 3 biological replicates. **d**, Representative confocal images showing expression of SHOX2, TBX18, and MYH6 following treatment with the indicated drugs. Scale bar=20 µm. **e**, Quantification of the percentage of SHOX2⁺TBX18⁺ cells in control-, forskolin-, and XMU-MP-1–treated cells. ***p < 0.001; *p < 0.05. Data are shown as mean ± SEM N = 3 biological replicates. **f**, qRT–PCR analysis of *HCN4* and *ISL1* expression in day 20 cells following drug treatment. Data are shown as mean ± SEM; ****p < 0.0001. N = 4 biological replicates.

**Extended Data Figure 14.**
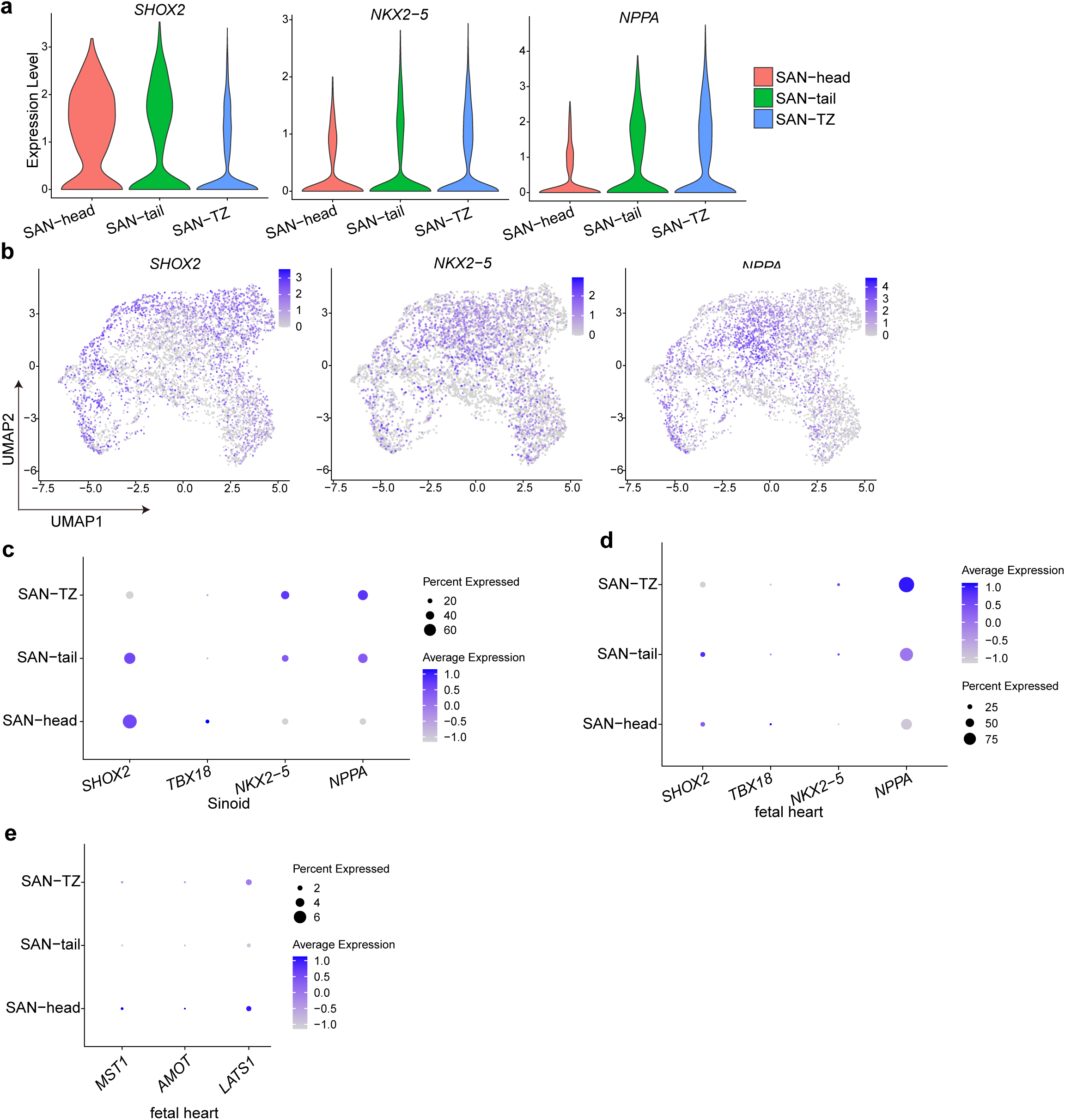
Subclustering analysis of SAN cells in hESC-derived Sinoids following Hippo pathway modulation and in human fetal SAN tissue. **a,** Violin plot displaying the expression of representative markers across SAN-head, SAN-tail, and SAN-TZ subclusters. **b,** UMAP displaying the expression of representative markers across SAN-head, SAN-tail, and SAN-TZ subclusters. **c,** Dot plot displaying the expression of representative markers across SAN-head, SAN-tail, and SAN-TZ subclusters. **d,** Dot plot displaying the expression of representative markers across SAN-head, SAN-tail, and SAN-TZ subclusters of human fetal SAN tissue. **e,** Dot plot displaying Hippo pathway associated genes in SAN-head, SAN-tail, and SAN-TZ subclusters of human fetal SAN tissue.

**Extended Data Figure 15.**
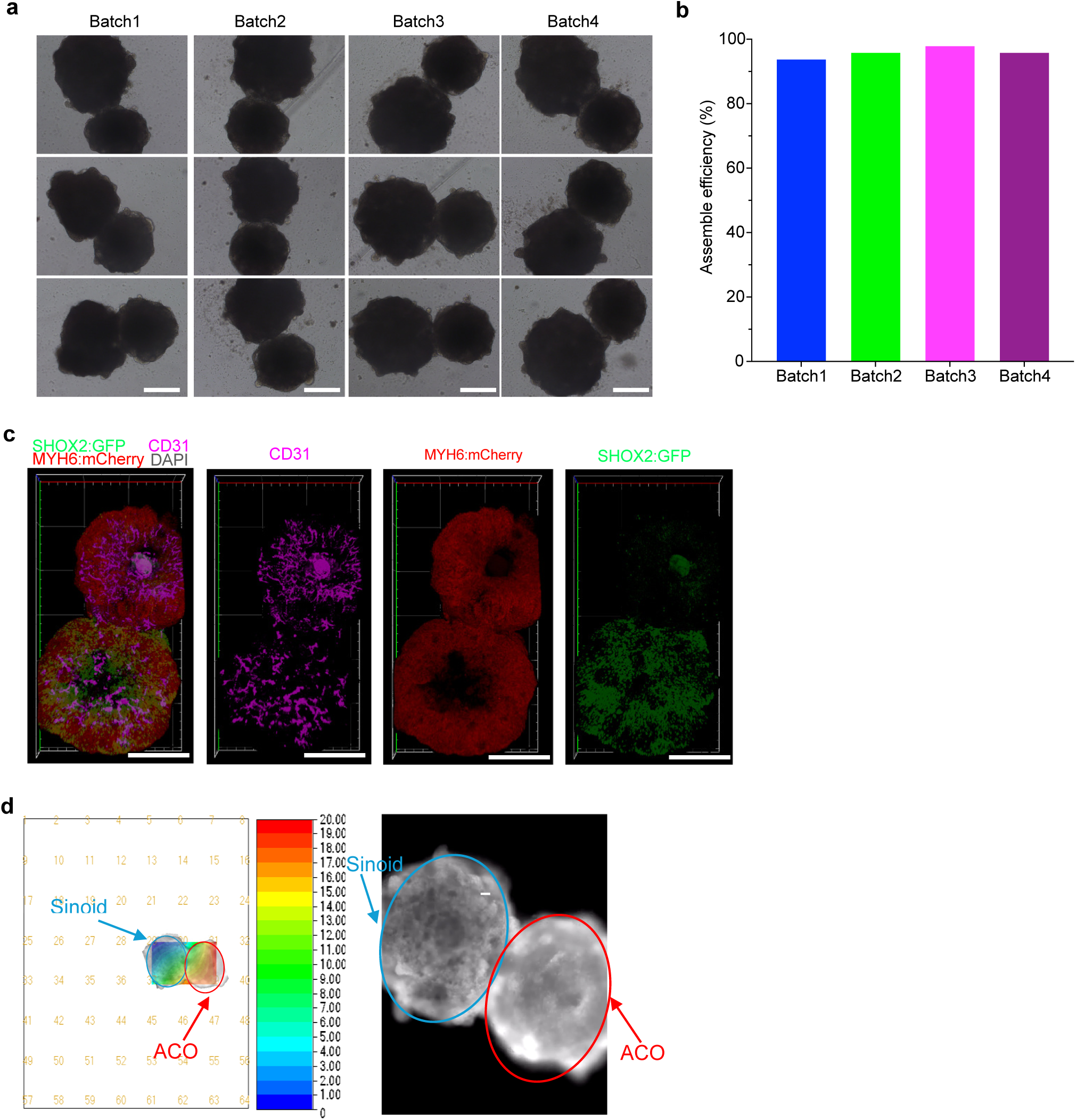
Characterization of hESC-derived SAN-PACOs. **a,** Bright field images of hESC-derived SAN-PACOs across multiple batches of differentiation. Scale bars=100 μm. **b,** Quantification of assembling efficiency of hESC-derived SAN-PACOs across multiple batches of differentiation. **c,** Representative confocal images of day 20 hPSC-derived SAN-PACOs showing expression of CD31. Scale bars=100 μm. **d,** Conduction velocity map from MEA analysis of an hESC-derived SAN-PACO.

**Extended Data Figure 16.**
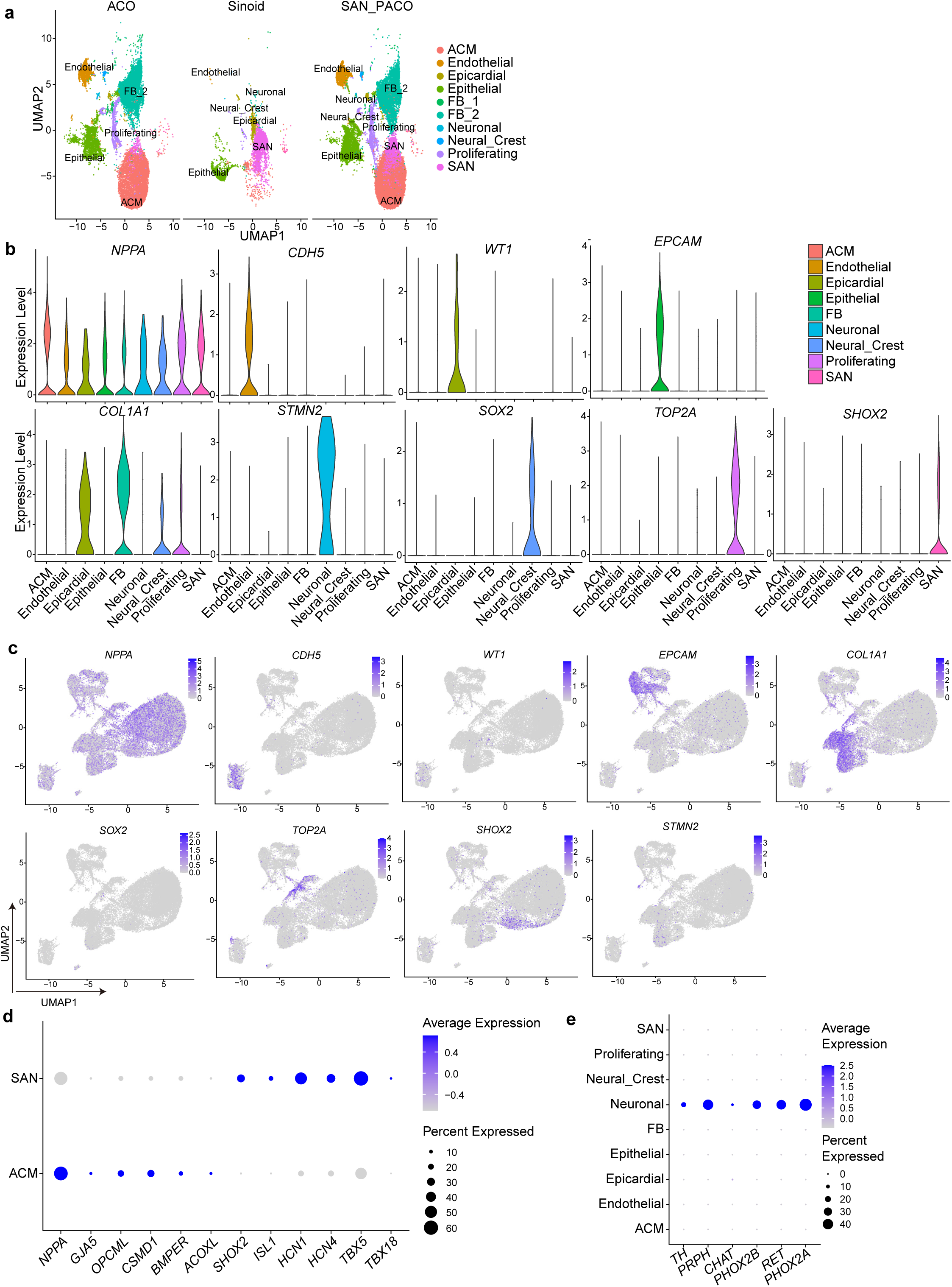
scRNA-seq analysis of SAN-PACO. **a,** Individual UMAP plots of hESC-derived Sinoids, ACOs, and SAN-PACO. **b,** Violin plots illustrating cell type–specific markers in SAN-PACOs. **c,** UMAP plots illustrating cell type–specific markers in SAN-PACOs. **d,** Dot plot illustrating expression of SAN and ACM marker genes within the SAN and ACM clusters of hESC-derived SAN-PACOs. **e,** Dot plot illustrating expression of AN genes within SAN-PACOs.

**Extended Data Figure 17.**
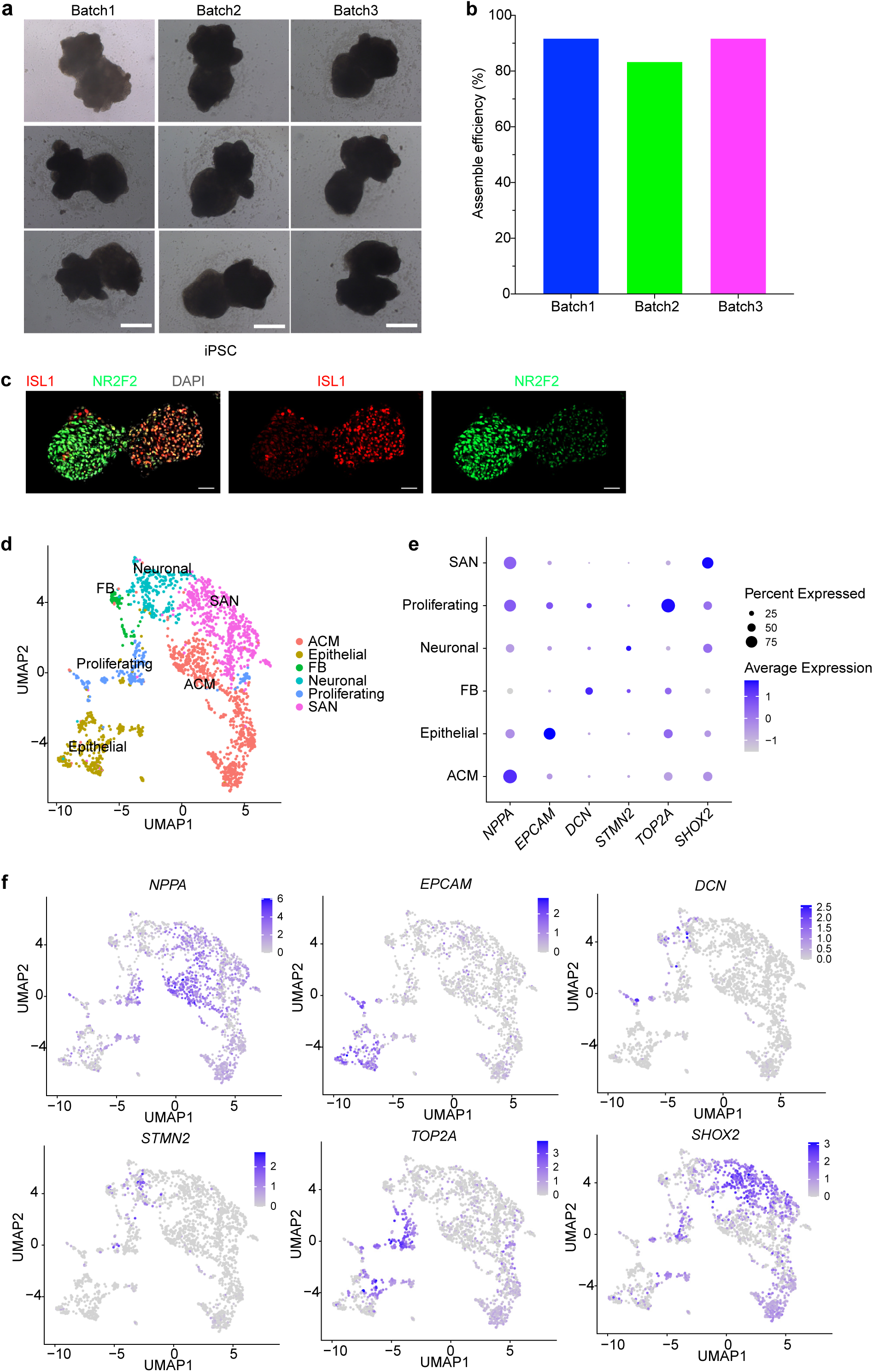
Characterization of hiPSC-derived SAN-PACOs. **a,** Bright field images of hiPSC-derived SAN-PACOs across multiple batches of differentiation. Scale bars=100 μm. **b,** Quantification of assembling efficiency of hiPSC-derived SAN-PACOs across multiple batches of differentiation. **c,** Representative confocal images of day 20 hiPSC-derived SAN-PACOs showing expression of ISL1 and NR2F2. Scale bars=50 μm. **d,** UMAP plot of hiPSC-derived SAN-PACOs. **e,** Dot plot of hiPSC-derived SAN-PACOs. **f,** UMAP plots displaying the expression patterns of selected marker genes in hiPSC-derived SAN-PACOs.

**Extended Data Figure 18.**
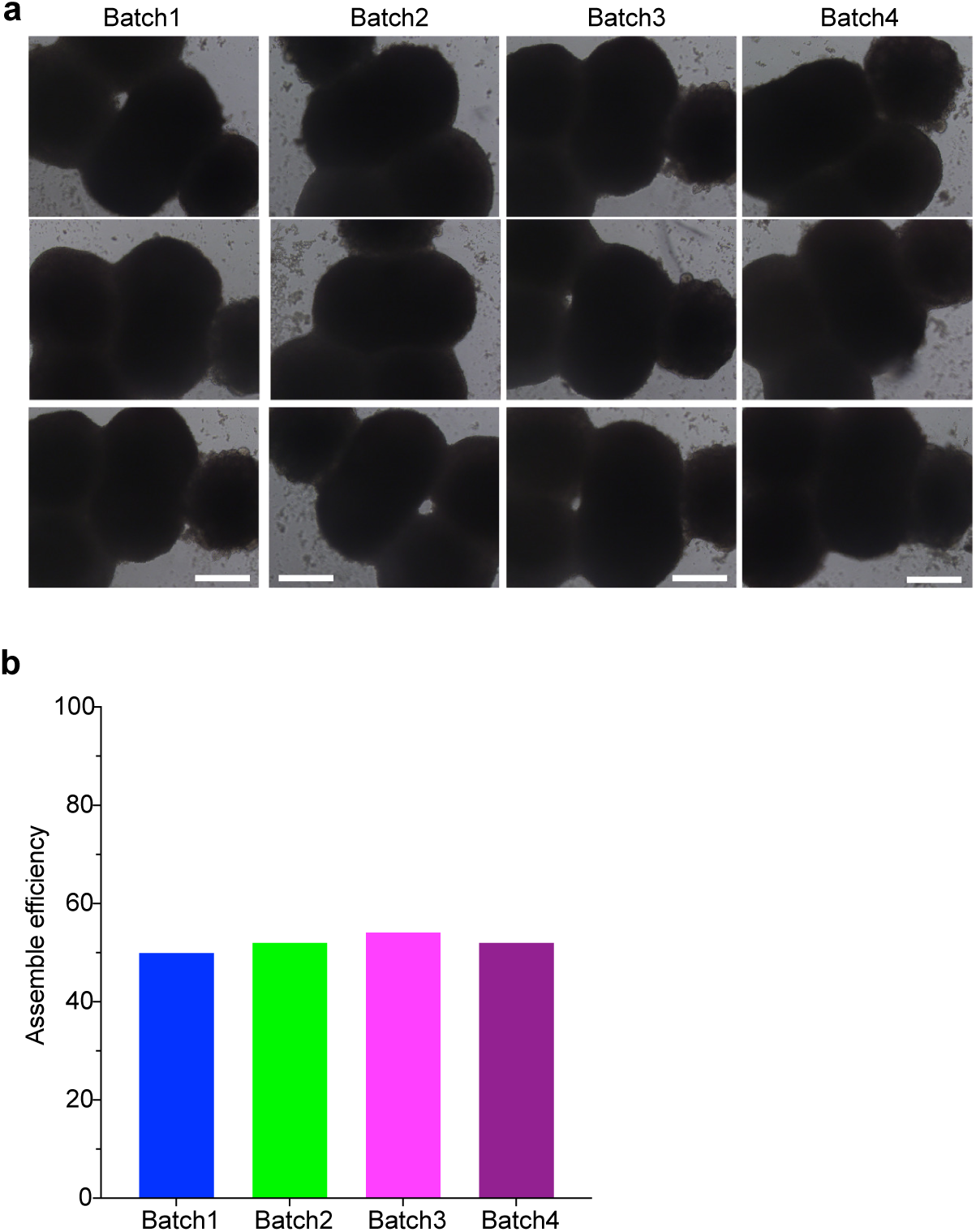
Characterization of mini-heart organoids. **a,** Bright field images of hESC-derived mini-heart organoids across multiple batches of differentiation. Scale bars=100 μm. **b,** Quantification of assembling efficiency of mini-heart organoids.

**Extended Data Figure 19.**
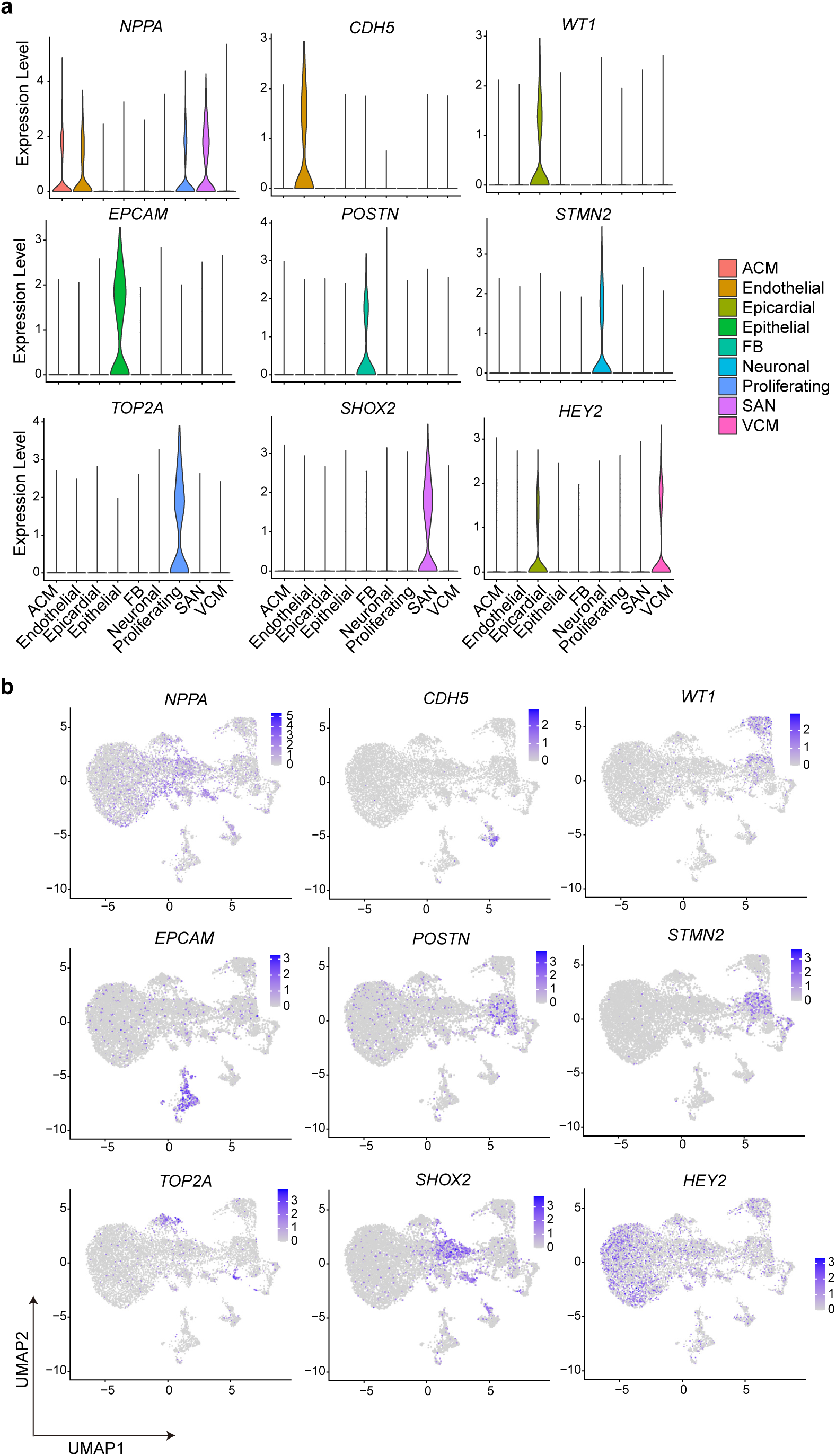
scRNA-seq analysis of hESC-derived mini-heart organoids. **a,** Violin plots displaying cell type–specific markers in hESC-derived mini-heart organoids. **b,** UMAP plots illustrating cell type–specific markers in hESC-derived mini-heart organoids.

## References.

1 Kennedy, A., et al. The Cardiac Conduction System: Generation and Conduction of the Cardiac Impulse. Crit Care Nurs Clin North Am 28, 269–279 (2016). 10.1016/j.cnc.2016.04.001

2 van Weerd, J. H. & Christoffels, V. M. The formation and function of the cardiac conduction system. Development 143, 197–210 (2016). 10.1242/dev.124883

3 de Soysa, T. Y., et al. Single-cell analysis of cardiogenesis reveals basis for organ-level developmental defects. Nature 572, 120–124 (2019). 10.1038/s41586-019-1414-x

4 Jia, G., et al. Single cell RNA-seq and ATAC-seq analysis of cardiac progenitor cell transition states and lineage settlement. Nat Commun 9, 4877 (2018). 10.1038/s41467-018-07307-6

5 Xiong, H., et al. Single-Cell Transcriptomics Reveals Chemotaxis-Mediated Intraorgan Crosstalk During Cardiogenesis. Circ Res 125, 398–410 (2019). 10.1161/CIRCRESAHA.119.315243

6 Goodyer, W. R., et al. Transcriptomic Profiling of the Developing Cardiac Conduction System at Single-Cell Resolution. Circ Res 125, 379–397 (2019). 10.1161/circresaha.118.314578

7 Hill, M. C., et al. A cellular atlas of Pitx2-dependent cardiac development. Development 146 (2019). 10.1242/dev.180398

8 Farah, E. N., et al. Spatially organized cellular communities form the developing human heart. Nature 627, 854–864 (2024). 10.1038/s41586-024-07171-z

9 Lim, A. A., et al. Single-cell transcriptome analysis reveals CD34 as a marker of human sinoatrial node pacemaker cardiomyocytes. Nat Commun 15, 10206 (2024). 10.1038/s41467-024-54337-4

10 Hou, X., et al. Single-cell RNA sequencing reveals the gene expression profile and cellular communication in human fetal heart development. Dev Biol 514, 87–98 (2024). 10.1016/j.ydbio.2024.06.004

11 Du, L., et al. Single cell and lineage tracing studies reveal the impact of CD34(+) cells on myocardial fibrosis during heart failure. Stem Cell Res Ther 14, 33 (2023). 10.1186/s13287-023-03256-0

12 Hocker, J. D., et al. Cardiac cell type-specific gene regulatory programs and disease risk association. Sci Adv 7 (2021). 10.1126/sciadv.abf1444

13 Rao, M., et al. Resolving the intertwining of inflammation and fibrosis in human heart failure at single-cell level. Basic Res Cardiol 116, 55 (2021). 10.1007/s00395-021-00897-1

14 Sim, C. B., et al. Sex-Specific Control of Human Heart Maturation by the Progesterone Receptor. Circulation 143, 1614–1628 (2021). 10.1161/circulationaha.120.051921

15 Selewa, A., et al. Single-cell genomics improves the discovery of risk variants and genes of atrial fibrillation. Nat Commun 14, 4999 (2023). 10.1038/s41467-023-40505-5

16 Hulsmans, M., et al. Recruited macrophages elicit atrial fibrillation. Science 381, 231–239 (2023). 10.1126/science.abq3061

17 Asp, M., et al. A Spatiotemporal Organ-Wide Gene Expression and Cell Atlas of the Developing Human Heart. Cell 179, 1647–1660 e1619 (2019). 10.1016/j.cell.2019.11.025

18 Cui, Y., et al. Single-Cell Transcriptome Analysis Maps the Developmental Track of the Human Heart. Cell Rep 26, 1934–1950 e1935 (2019). 10.1016/j.celrep.2019.01.079

19 Litvinukova, M., et al. Cells of the adult human heart. Nature 588, 466–472 (2020). 10.1038/s41586-020-2797-4

20 Marin-Sedeno, E., de Morentin, X. M., Perez-Pomares, J. M., Gomez-Cabrero, D. & Ruiz-Villalba, A. Understanding the Adult Mammalian Heart at Single-Cell RNA-Seq Resolution. Front Cell Dev Biol 9, 645276 (2021). 10.3389/fcell.2021.645276

21 Tabula Muris, C., et al. Single-cell transcriptomics of 20 mouse organs creates a Tabula Muris. Nature 562, 367–372 (2018). 10.1038/s41586-018-0590-4

22 Verheijck, E. E., et al. Electrophysiological features of the mouse sinoatrial node in relation to connexin distribution. Cardiovasc Res 52, 40–50 (2001). 10.1016/s0008-6363(01)00364-9

23 Kanemaru, K., et al. Spatially resolved multiomics of human cardiac niches. Nature 619, 801–810 (2023). 10.1038/s41586-023-06311-1

24 Asp, M., et al. A Spatiotemporal Organ-Wide Gene Expression and Cell Atlas of the Developing Human Heart. Cell 179, 1647–1660.e1619 (2019). 10.1016/j.cell.2019.11.025

25 Schep, A. N., Wu, B., Buenrostro, J. D. & Greenleaf, W. J. chromVAR: inferring transcription-factor-associated accessibility from single-cell epigenomic data. Nat Methods 14, 975–978 (2017). 10.1038/nmeth.4401

26 Bosada, F. M., et al. An atrial fibrillation-associated regulatory region modulates cardiac Tbx5 levels and arrhythmia susceptibility. Elife 12 (2023). 10.7554/eLife.80317

27 Weng, L. C., et al. The impact of common and rare genetic variants on bradyarrhythmia development. Nat Genet 57, 53–64 (2025). 10.1038/s41588-024-01978-2

28 Song, M., et al. GATA4/5/6 family transcription factors are conserved determinants of cardiac versus pharyngeal mesoderm fate. Sci Adv 8, eabg0834 (2022). 10.1126/sciadv.abg0834

29 Gharibeh, L., et al. GATA6 is a regulator of sinus node development and heart rhythm. Proc. Natl. Acad. Sci. U. S. A. 118 (2021). 10.1073/pnas.2007322118

30 Lin, Z., et al. Evolutionary-scale prediction of atomic-level protein structure with a language model. Science 379, 1123–1130 (2023). 10.1126/science.ade2574

31 Nadig, A., et al. Transcriptome-wide analysis of differential expression in perturbation atlases. Nat Genet 57, 1228–1237 (2025). 10.1038/s41588-025-02169-3

32 Roohani, Y. H., et al. Virtual Cell Challenge: Toward a Turing test for the virtual cell. Cell 188, 3370–3374 (2025). 10.1016/j.cell.2025.06.008

33 Kim, K. H., Nakaoka, Y., Augustin, H. G. & Koh, G. Y. Myocardial Angiopoietin-1 Controls Atrial Chamber Morphogenesis by Spatiotemporal Degradation of Cardiac Jelly. Cell Rep 23, 2455–2466 (2018). 10.1016/j.celrep.2018.04.080

34 Manzo, A., et al. CCL21 expression pattern of human secondary lymphoid organ stroma is conserved in inflammatory lesions with lymphoid neogenesis. Am J Pathol 171, 1549–1562 (2007). 10.2353/ajpath.2007.061275

35 van der Maarel, L. E., Postma, A. V. & Christoffels, V. M. Genetics of sinoatrial node function and heart rate disorders. Dis Model Mech 16 (2023). 10.1242/dmm.050101

36 Vedantham, V., Galang, G., Evangelista, M., Deo, R. C. & Srivastava, D. RNA sequencing of mouse sinoatrial node reveals an upstream regulatory role for Islet-1 in cardiac pacemaker cells. Circ Res 116, 797–803 (2015). 10.1161/circresaha.116.305913

37 Marionneau, C., et al. Specific pattern of ionic channel gene expression associated with pacemaker activity in the mouse heart. J Physiol 562, 223–234 (2005). 10.1113/jphysiol.2004.074047

38 Takayama, Y., et al. Selective Induction of Human Autonomic Neurons Enables Precise Control of Cardiomyocyte Beating. Sci Rep 10, 9464 (2020). 10.1038/s41598-020-66303-3

39 Lamar, J. M., et al. The Hippo pathway target, YAP, promotes metastasis through its TEAD-interaction domain. Proc Natl Acad Sci U S A 109, E2441–2450 (2012). 10.1073/pnas.1212021109

40 Yuan, Y., et al. YAP1/TAZ-TEAD transcriptional networks maintain skin homeostasis by regulating cell proliferation and limiting KLF4 activity. Nat Commun 11, 1472 (2020). 10.1038/s41467-020-15301-0

41 Ghazizadeh, Z., et al. A dual SHOX2:GFP; MYH6:mCherry knockin hESC reporter line for derivation of human SAN-like cells. iScience 25, 104153 (2022). 10.1016/j.isci.2022.104153

42 Han, Y., et al. SARS-CoV-2 Infection Induces Ferroptosis of Sinoatrial Node Pacemaker Cells. Circ Res 130, 963–977 (2022). 10.1161/circresaha.121.320518

43 Tran, K., et al. Identification of small molecules that protect pancreatic β cells against endoplasmic reticulum stress-induced cell death. ACS Chem Biol 9, 2796–2806 (2014). 10.1021/cb500740d

44 Li, W. et al. Nanchangmycin regulates FYN, PTK2, and MAPK1/3 to control the fibrotic activity of human hepatic stellate cells. Elife 11 (2022). 10.7554/eLife.74513

45 Paffett-Lugassy, N., et al. Heart field origin of great vessel precursors relies on nkx2.5-mediated vasculogenesis. Nat Cell Biol 15, 1362–1369 (2013). 10.1038/ncb2862

46 Colombo, S., et al. Nkx genes establish second heart field cardiomyocyte progenitors at the arterial pole and pattern the venous pole through Isl1 repression. Development 145 (2018). 10.1242/dev.161497

47 Graham, K., et al. Discovery of YAP1/TAZ pathway inhibitors through phenotypic screening with potent anti-tumor activity via blockade of Rho-GTPase signaling. Cell Chem Biol 31, 1247–1263.e1216 (2024). 10.1016/j.chembiol.2024.02.013

48 Niederreither, K., et al. Embryonic retinoic acid synthesis is essential for heart morphogenesis in the mouse. Development 128, 1019–1031 (2001). 10.1242/dev.128.7.1019

49 Zhang, Q., et al. Direct differentiation of atrial and ventricular myocytes from human embryonic stem cells by alternating retinoid signals. Cell Res 21, 579–587 (2011). 10.1038/cr.2010.163

50 Li, J., et al. Modeling the atrioventricular conduction axis using human pluripotent stem cell-derived cardiac assembloids. Cell Stem Cell 31, 1667–1684.e1666 (2024). 10.1016/j.stem.2024.08.008

51 Boyett, M. R., Honjo, H. & Kodama, I. The sinoatrial node, a heterogeneous pacemaker structure. Cardiovascular Research 47, 658–687 (2000). 10.1016/s0008-6363(00)00135-8

52 Wilhelm, T. I., et al. Electroanatomical Conduction Characteristics of Pig Myocardial Tissue Derived from High-Density Mapping. J Clin Med 12 (2023). 10.3390/jcm12175598

53 Lim, A. A., et al. Single-cell transcriptome analysis reveals CD34 as a marker of human sinoatrial node pacemaker cardiomyocytes. Nature Communications 15, 10206 (2024). 10.1038/s41467-024-54337-4

54 Yao, Z., et al. A high-resolution transcriptomic and spatial atlas of cell types in the whole mouse brain. Nature 624, 317–332 (2023). 10.1038/s41586-023-06812-z

55 Siletti, K., et al. Transcriptomic diversity of cell types across the adult human brain. Science 382, eadd7046 (2023). 10.1126/science.add7046

56 Wiesinger, A., et al. A single cell transcriptional roadmap of human pacemaker cell differentiation. Elife 11 (2022). 10.7554/eLife.76781

57 Wiesinger, A., et al. A single cell transcriptional roadmap of human pacemaker cell differentiation. eLife 11, e76781 (2022). 10.7554/eLife.76781

58 Kim, M., et al. cAMP/PKA signalling reinforces the LATS-YAP pathway to fully suppress YAP in response to actin cytoskeletal changes. Embo j 32, 1543–1555 (2013). 10.1038/emboj.2013.102

59 Mitchell, E., Mellor, C. E. L. & Purba, T. S. XMU-MP-1 induces growth arrest in a model human mini-organ and antagonises cell cycle-dependent paclitaxel cytotoxicity. Cell Div 15, 11 (2020). 10.1186/s13008-020-00067-0

60 Oh, Y., et al. Functional Coupling with Cardiac Muscle Promotes Maturation of hPSC-Derived Sympathetic Neurons. Cell Stem Cell 19, 95–106 (2016). 10.1016/j.stem.2016.05.002

61 Protze, S. I., et al. Sinoatrial node cardiomyocytes derived from human pluripotent cells function as a biological pacemaker. Nature Biotechnology 35, 56–68 (2017). 10.1038/nbt.3745

62 Yechikov, S., et al. NODAL inhibition promotes differentiation of pacemaker-like cardiomyocytes from human induced pluripotent stem cells. Stem Cell Res 49, 102043 (2020). 10.1016/j.scr.2020.102043

63 Liang, W., et al. Canonical Wnt signaling promotes pacemaker cell specification of cardiac mesodermal cells derived from mouse and human embryonic stem cells. Stem Cells 38, 352–368 (2020). 10.1002/stem.3106

64 Ren, J., et al. Canonical Wnt5b Signaling Directs Outlying Nkx2.5+ Mesoderm into Pacemaker Cardiomyocytes. Dev Cell 50, 729–743.e725 (2019). 10.1016/j.devcel.2019.07.014

65 Zhao, H., et al. Overexpression of TBX3 in human induced pluripotent stem cells (hiPSCs) increases their differentiation into cardiac pacemaker-like cells. Biomed Pharmacother 130, 110612 (2020). 10.1016/j.biopha.2020.110612

66 Gorabi, A. M., et al. TBX18 transcription factor overexpression in human-induced pluripotent stem cells increases their differentiation into pacemaker-like cells. J Cell Physiol 234, 1534–1546 (2019). 10.1002/jcp.27018

67 Darche, F. F., et al. Improved Generation of Human Induced Pluripotent Stem Cell-Derived Cardiac Pacemaker Cells Using Novel Differentiation Protocols. Int J Mol Sci 23 (2022). 10.3390/ijms23137318

68 Li, J., et al. Molecular and electrophysiological evaluation of human cardiomyocyte subtypes to facilitate generation of composite cardiac models. J Tissue Eng 13, 20417314221127908 (2022). 10.1177/20417314221127908

69 Xin, M., et al. Hippo pathway effector Yap promotes cardiac regeneration. Proc. Natl. Acad. Sci. U. S. A. 110, 13839–13844 (2013). 10.1073/pnas.1313192110

70 Vinarsky, V., et al. YAP1 reactivation in cardiomyocytes following ECM remodelling contributes to the development of contractile force and sarcomere maturation. Cell Death Discov 11, 518 (2025). 10.1038/s41420-025-02793-2

71 Singh, V. P., et al. Hippo Pathway Effector Tead1 Induces Cardiac Fibroblast to Cardiomyocyte Reprogramming. Journal of the American Heart Association 10, e022659 (2021). doi:10.1161/JAHA.121.022659

72 Hao, J., Galindo, Cristi L., Tran, T.-L. & Sawyer, Douglas B. Neuregulin-1β induces embryonic stem cell cardiomyogenesis via ErbB3/ErbB2 receptors. Biochemical Journal 458, 335–341 (2014). 10.1042/bj20130818

73 Zhu, W. Z., et al. Neuregulin/ErbB signaling regulates cardiac subtype specification in differentiating human embryonic stem cells. Circ Res 107, 776–786 (2010). 10.1161/circresaha.110.223917

74 Miao, Y., et al. Co-development of mesoderm and endoderm enables organotypic vascularization in lung and gut organoids. Cell 188, 4295–4313.e4227 (2025). 10.1016/j.cell.2025.05.041

75 Bouffi, C., et al. In vivo development of immune tissue in human intestinal organoids transplanted into humanized mice. Nat Biotechnol 41, 824–831 (2023). 10.1038/s41587-022-01558-x

76 Schmidt, C., et al. Multi-chamber cardioids unravel human heart development and cardiac defects. Cell 186, 5587–5605.e5527 (2023). 10.1016/j.cell.2023.10.030

77 Hao, Y., et al. Integrated analysis of multimodal single-cell data. Cell 184, 3573–3587.e3529 (2021). 10.1016/j.cell.2021.04.048

78 Luo, W., Friedman, M. S., Shedden, K., Hankenson, K. D. & Woolf, P. J. GAGE: generally applicable gene set enrichment for pathway analysis. BMC Bioinformatics 10, 161 (2009). 10.1186/1471-2105-10-161

79 Paffett-Lugassy, N., et al. Unique developmental trajectories and genetic regulation of ventricular and outflow tract progenitors in the zebrafish second heart field. Development 144, 4616–4624 (2017). 10.1242/dev.153411

80 Huang, Y., et al. Building a literature knowledge base towards transparent biomedical AI. bioRxiv, 2024.2009.2022.614323 (2025). 10.1101/2024.09.22.614323

